# Auxin Exposure Disrupts Feeding Behavior and Fatty Acid Metabolism in Adult *Drosophila*

**DOI:** 10.1101/2023.08.15.553385

**Authors:** Sophie A. Fleck, Puja Biswas, Emily D. DeWitt, Rebecca L. Knuteson, Robert C. Eisman, Travis Nemkov, Angelo D’Alessandro, Jason M. Tennessen, Elizabeth J. Rideout, Lesley N. Weaver

## Abstract

The ease of genetic manipulation in *Drosophila melanogaster* using the *Gal4/UAS* system has been beneficial in addressing key biological questions. Current modifications of this methodology to temporally induce transgene expression require temperature changes or exposure to exogenous compounds, both of which have been shown to have detrimental effects on physiological processes. The recently described auxin-inducible gene expression system (AGES) utilizes the plant hormone auxin to induce transgene expression and is proposed to be the least toxic compound for genetic manipulation, with no obvious effects on *Drosophila* development and survival in one wild-type strain. Here we show that auxin delays larval development in another widely-used fly strain, and that short- and long-term auxin exposure in adult *Drosophila* induces observable changes in physiology and feeding behavior. We further reveal a dosage response to adult survival upon auxin exposure, and that the recommended auxin concentration for AGES alters feeding activity. Furthermore, auxin fed male and female flies exhibit a significant decrease in triglyceride levels and display altered transcription of fatty acid metabolism genes. Although fatty acid metabolism is disrupted, auxin does not significantly impact adult female fecundity or progeny survival, suggesting AGES may be an ideal methodology for studying limited biological processes. These results emphasize that experiments using temporal binary systems must be carefully designed and controlled to avoid confounding effects and misinterpretation of results.

## INTRODUCTION

The intricate dissection of cell-type specific processes in *Drosophila* is largely dependent on the yeast-derived *Gal4/UAS* binary system, which allows manipulation of biological pathways in a spatial and temporal manner (BRAND AND PERRIMON 1993). This methodology utilizes the Gal4 transcription factor that is under control of a tissue-specific promoter to induce transgene expression downstream of an <u>U</u>pstream <u>A</u>ctivating <u>S</u>equence (UAS). Conditional control of gene expression using changes in temperature (MCGUIRE *et al*. 2003) or feeding of small molecules (ROMAN *et al*. 2001; POTTER *et al*. 2010; MCCLURE *et al*. 2022) restricts Gal4 activity to specific developmental timepoints. For example, use of a temperature sensitive *Gal80* mutant transgene [*Gal80^ts^*, an inhibitor of Gal4 (DOUGLAS AND HAWTHORNE 1966)] allows for temporal control of Gal4 activity with a simple shift to the *Gal80^ts^* restrictive temperature [29°C, (MCGUIRE *et al*. 2003)]. Conversely, drug-inducible systems such as *GeneSwitch* and the *Q-system* control temporal and reversible transgene expression with administration of RU486 or quinic acid, respectively, without the need to rear flies at the *Gal80^ts^* permissive temperature (18°C). These modifications to the *Gal4/UAS* system have improved the capability to characterize the roles of essential biological pathways in a tissue-specific manner while avoiding lethality at key developmental stages.

Despite these advancements, each methodology has caveats that must be considered. For example, rearing flies containing *Gal80^ts^* at 18°C nearly doubles the developmental time from egg to adult (POWSNER 1935). In addition, the relatively high restrictive temperature needed to inactivate Gal80^ts^ has adverse effects on physiological processes including circadian rhythm (PARISKY *et al*. 2016), aging (MIQUEL *et al*. 1976), and progeny survival (GANDARA AND DRUMMOND-BARBOSA 2022). Similarly, use of RU486 has been demonstrated to repress muscle-specific mitochondrial genes (ROBLES-MURGUIA *et al*. 2019) and lipogenesis (MA *et al*. 2021), among other defects (LANDIS *et al*. 2015; YAMADA *et al*. 2017), making the *GeneSwitch* system non-ideal for certain experiments. Although designed to provide flexibility in experimental design and remove temperature-related defects, the alterations in physiology, behavior, and lifespan [some of which are not shared between the sexes (LANDIS *et al*. 2015)] due to RU486 feeding impose difficulties in data interpretation.

The <u>a</u>uxin-inducible <u>g</u>ene <u>e</u>xpression <u>s</u>ystem (AGES) was recently developed as an alternative method to induce transgene expression and is compatible with the breadth of *Gal4* transgenic lines available (MCCLURE *et al*. 2022). In this system, *Gal80* is fused to auxin-inducible degron tags that target Gal80 for degradation, allowing for Gal4-mediated transgene induction upon auxin consumption. This system poses substantial advantages. For example, flies can be reared at the optimal temperature for development (25°C) and transgene expression is strictly induced with supplementation of auxin to the media. In addition, both control and experimental animals are genetically identical (similarly to the *GeneSwitch* and *Q-system*), thus minimizing differences that may arise due to genetic variation.

However, there is emerging evidence that insects can synthesize auxin (YOKOYAMA *et al*. 2017; TOKUDA *et al*. 2022), suggesting that there may be important biological roles for this hormone in *Drosophila*. While exposure to 10 mM 1-naphthaleneacetic acid (the widely employed synthetic hormone in the auxin family and hereafter referred to as “auxin”) has been reported to have no effect on insect development, survival, or movement in a wild-type *Drosophila* strain (MCCLURE *et al*. 2022), it remains unclear whether auxin exposure affects development in other commonly used *Drosophila* genetic background strains. It is also unknown whether increased auxin exposure in adult *Drosophila* results in subtle defects in physiological processes that may confound experimental interpretations. In this study we sought to test whether auxin affects larval development in additional strains, and to determine whether auxin exposure in adults leads to defects in metabolism, the transcriptome, and oogenesis. We found that recommended concentrations of auxin for AGES disrupt feeding behavior, whereas increasing levels of auxin results in physiological changes and lethality. Additionally, we found that auxin exposure delays larval development, decreases triglyceride levels and alters the transcriptomic profile of fatty acid metabolism genes in both sexes. Finally, despite the decrease in circulating lipids, auxin does not severely disrupt processes of oogenesis or progeny survival, suggesting AGES may be an appropriate method to use for some studies. Our results highlight changes in development and physiology that should be considered when using auxin to manipulate gene and protein expression in larval and adult *Drosophila,* as well as other factors researchers should account for in experimental design using temporal control of the *Gal4/UAS* system.

## MATERIALS AND METHODS

### *Drosophila* strains and culture

*Drosophila* stocks were maintained on Bloomington *Drosophila* Stock Center (BDSC) Cornmeal Food that consists of 15.9 g/L inactive yeast, 9.2 g/L soy flour, 67 g/L yellow cornmeal, 5.3 g/L agar, 70.6 g/L light corn syrup, 0.059 M propionic acid at 22-25°C. Medium was supplemented with inactive wet yeast paste for all experiments, unless otherwise noted. For a subset of experiments, where indicated, flies were reared on yeast-sugar-cornmeal food (LEWIS 1960) that consists of 20.5 g/L white sugar, 70.9 g/L D-glucose, 48.5 g/L cornmeal, 30.3 g/L yeast, 0.5 g/L CaCl_2_-2H_2_O, 0.5 g/L MgSO_4_-7H_2_O, 4.6 g/L agar, 4.9 ml/L propionic acid, 0.49 ml/L phosphoric acid. For experiments using this diet, the food was not supplemented with inactive wet yeast paste. The previously described *w^1118^; tubP-TIR1-T2A-Gal80.AID* [“*AGES*”, (MCCLURE *et al*. 2022)] line was used and obtained from the Bloomington *Drosophila* Stock Center (BDSC 92470; https://bdsc.indiana.edu). *Oregon-R* (BDSC 25211) and *y^1^ w^1118^; VK00040/TM6B* (BDSC 9755) lines were used as controls. Flies from the *w^1118^* strain (BDSC 3605) were used to monitor larval development and adult triglyceride levels. Balancer chromosomes and other genetic elements are described in Flybase (www.flybase.org).

For all ovarian experiments, 0-to-2-day-old females were mated with *AGES* males and incubated at 25°C for up to 15 days at ≥70% humidity on medium containing either 1 mM NaOH [0 mM 1-naphthaleneacetic acid (referred to as “auxin” throughout the manuscript)] or 10 mM auxin dissolved in 1 mM NaOH [the recommended concentration of auxin (MCCLURE *et al*. 2022)]. Medium was supplemented with inactive wet yeast containing either 1 mM NaOH or 10 mM auxin daily, except where noted.

### Larval development time

*w^1118^* females laid eggs on grape plates supplemented with yeast paste during a 3-hr collection period. Newly-hatched larvae were transferred to the yeast-sugar-cornmeal diet at a density of 50 larvae per 10 ml food. For the auxin-containing medium, auxin was added to the recommended concentration of 5 mM to cooled fly food immediately prior to filling vials (MCCLURE *et al*. 2022). Percent pupation was calculated by comparing the number of pupae at each 12 hr interval to the total pupae in the vial.

### Auxin exposure dose response curves

Dose response curves were performed as previously described (HOLSOPPLE *et al*. 2023). Briefly, zero-to-two-day old flies were transferred to bottles containing fresh BDSC food and aged for two days. Flies were sorted by sex in groups of 20 per vial and aged for an additional 48 hours at 25°C to allow recovery from CO_2_ anesthesia. Flies were then transferred to starvation vials containing sterile milli-Q water for 16 hours at 25°C. Following starvation, flies were transferred to exposure vials containing liquid food [4% sucrose (m/v), 1.5% yeast extract (m/v), 1 mM NaOH] and 0, 2, 4, 6, 10, 15, 20, 25, 90, 125, or 350 mM auxin. The number of dead female or male flies per vial were counted in six replicates per concentration and the percentage of dead flies was subjected to mathematical modeling through the Benchmark Dose Software (BMDS) published by the U.S. Environmental Protection Agency (EPA; https://www.epa.gov/bmds).

### Fly Liquid-Food Interaction Counter (FLIC) assay

The Fly Liquid-Food Interaction Counter (FLIC) system was used to determine differences in feeding behavior in male and female flies exposed to food with or without auxin as previously described (RO *et al*. 2014). FLIC *Drosophila* Feeding Monitors (DFMs, Sable Systems International, models DFMV2 and DFMV3) were used in the single choice configuration and each chamber was loaded with 0 mM auxin liquid food solution [4% sucrose (m/v), 1.5% yeast extract (m/v), 1 mM NaOH] or the recommended dose of 10 mM auxin liquid food [4% sucrose (m/v), 1.5% yeast extract (m/v), 10 mM auxin]. Four-to-six-day old flies were briefly anesthetized and aspirated into the DFM chambers. Feeding behavior was measured for 24 hours. Each FLIC experiment contains pooled data from at least 30 flies for each genotype, sex, and auxin concentration condition. FLIC data were analyzed using previously described custom R code (RO *et al*. 2014), which is available at https://github.com/PletcherLab/FLIC_R_Code. Default thresholds were used for analysis except for the following: minimum feeding threshold = 10, tasting threshold = (0,10). Animals that did not participate (i.e., returned zero values), whose DFM returned an unstable baseline signal, or who produced extreme outliers (i.e., exceeding twice the mean of the population) were excluded from analysis. Data were subjected to a Mann-Whitney *U*-test.

### Ultra High-pressure Liquid Chromatography-Mass Spectrometry (UHPLC-MS)-based metabolomics and analysis

Adult male and female flies were exposed to 0 mM or the recommended dose of 10 mM auxin as described above for the dose response curve assays and were flash frozen in liquid nitrogen. Analyses were performed at the University of Colorado Anschutz Medical Campus, as previously described (NEMKOV *et al*. 2019; NEMKOV *et al*. 2022) with minor modifications. Briefly, the analytical platform employs a Vanquish UHPLC system (ThermoFisher Scientific) coupled online to a Q Exactive mass spectrometer (ThermoFisher Scientific). The (semi)polar extracts were resolved over a Kinetex C18 column, 2.1 x 30 mm, 1.7 µm particle size (Phenomenex) using a high-throughput 1 minute gradient. Solvents were supplemented with 0.1% formic acid for positive mode runs and 10 mM ammonium acetate +0.1% ammonium hydroxide for negative mode runs. The Q Exactive mass spectrometer (ThermoFisher Scientific) was operated independently in positive or negative ion mode, scanning in Full MS mode (2 μscans) from 60 to 900 m/z at 70,000 resolution, with 4 kV spray voltage, 45 sheath gas, 15 auxiliary gas. Calibration was performed prior to analysis using the Pierce^TM^ Positive and Negative Ion Calibration Solutions (ThermoFisher Scientific). Metabolomics raw data were processed using El-Maven (AGRAWAL *et al*. 2019) and analyzed using Metaboanalyst 5.0 (PANG *et al*. 2021), with the data first preprocessed using log normalization and Pareto scaling.

### Triglyceride assays

Zero-to-two-day old adult males and females were collected and maintained on BDSC food for 2 days. Two-to-four-day old adults were starved for 16 hours and then exposed to 0 mM or the recommended dose of 10 mM auxin liquid food for 48 hours as described above for the dose response assays. Whole bodies from five animals of each sex and genotype were washed in phosphate-buffered saline (PBS), pH 7.0 and flash frozen in liquid nitrogen. Samples were homogenized in 100 µl cold PBS + 0.05% Tween 20 (PBST) and heat-treated for 10 minutes at 90°C. The resulting homogenate was assayed for triglyceride (TAG) and soluble protein levels as previously described (TENNESSEN *et al*. 2014). TAG amounts were normalized to protein amounts and expressed as µg/ml TAG per µg/ml protein. Data were subjected to a paired Student’s *t*-test.

For TAG assays on *w^1118^* flies, newly hatched larvae were transferred to yeast-sugar-cornmeal food and reared at a density of 50 larvae per 10 ml food at 25°C. Male and female pupae were separated as late pupae according to sex combs. Two experimental designs were used. For one design, virgin male and female flies were kept at a density of 20 flies per 10 ml food (+/- recommended dose of 10 mM auxin) from eclosion until 5 days of age. Five-day-old male and female flies were collected, snap frozen at -80°C, weighed, and finally subjected to a TAG assay. In the second experimental design, newly-eclosed flies were subjected to one of three protocols: (1) maintained continuously on yeast-sugar-cornmeal food with no auxin, where flies were collected, frozen, weighed, and subjected to a TAG assay at 5 and 10 days of age; (2) maintained on yeast-sugar-cornmeal food with 10 mM auxin for 5 days and shifted to food with no auxin for a further 5 days, where flies were collected, frozen, weighed, and subjected to a TAG assay at 5 and 10 days of age; and (3) maintained continuously on yeast-sugar-cornmeal food with the recommended 10 mM dose of auxin, with flies collected, frozen, weighed, and subjected to a TAG assay at 5 and 10 days of age. Flies were flipped every two days in both experimental designs. One biological replicate of three or five flies was homogenized in 150 µl or 350 µl of 0.1% Tween in 1X phosphate-buffered saline (PBS) using 50 µl of glass beads agitated at 8 m/s for 5 sec. TAG assay was performed using the Stanbio Triglyceride Liquicolor kit (#SB2100430) according to manufacturer’s protocol. TAG is expressed as percent body fat as previously described (WAT *et al*. 2020). Each experiment includes four biological replicates, and each experiment was repeated twice for a total of eight biological replicates per sex; data were analyzed using either a Student’s *t*-test, one-way ANOVA, or two-way ANOVA, as indicated.

### RNA isolation, RNA sequencing, and data analysis

Twenty whole adult animals of each genotype and sex were exposed to 0 mM or the recommended dose of 10 mM auxin, as described above for the dose response curves, and flash frozen in liquid nitrogen. Tissue was lysed in 500 µl lysis buffer from the RNAqueous-4PCR DNA-free RNA isolation for RT-PCR kit (Ambion). RNA was extracted from all samples following manufacturer’s instructions. Three independent experiments were performed for RNA sequencing.

cDNA library construction, Illumina sequencing, and differential expression analysis was performed by Novogene Bioinformatics Technology Co., Ltd (Beijing, China). The cDNA libraries were prepared using the NEBNext Ultra RNA Library Prep Kit for Illumina (New England Biolabs) according to the manufacturer’s instructions. The cDNA library for each sample was quality assessed using an Agilent Bioanalyzer 2100, and library preparations were sequenced on a NovaSeq6000 platform with PE150 read lengths.

Reads obtained from sequencing were aligned to the *D. melanogaster* reference genome using the TopHat read alignment tool (TRAPNELL *et al*. 2009) for each of the sequencing datasets. The reference sequences were downloaded from the Ensembl project website (useast.ensembl.org). TopHat alignments were used to generate read counts for each gene using HTSeq (ANDERS *et al*. 2015), which were subsequently used to generate the differential expression results using the DESeq2 R package (ANDERS *et al*. 2015). RNA sequencing produced an average of 40,448,108 reads across the 36 sequencing libraries, ranging from 37,632,234 to 58,099,256 reads per sample (representing an average of 96.2% mapped to the *Drosophila* genome). Enriched genes with a corrected *P* value less than 0.05 were considered significant.

### cDNA synthesis and quantitative reverse-transcriptase polymerase chain reaction (qRT-PCR)

cDNA was synthesized from 500 ng of total RNA described above for each sample using Superscript II Reverse Transcriptase (Thermo Fisher Scientific) according to the manufacturer’s instructions. PowerUp SYBR Green Master Mix (Thermo Fisher Scientific) was used for RT-qPCR. The reactions for three independent biological replicates were performed in triplicate using LightCycler 96 (Roche). Amplification fluorescence threshold was determined by LightCycler 96 software, and ddCT were calculated using Microsoft Excel. Fold change of transcript levels was calculated in Excel as described (TAYLOR *et al*. 2019). The primers used for all PCR reactions are listed in Table S8. *Rp49* and *Act5C* transcript levels were used as references.

### Egg laying and hatching assays

Egg production was measured as previously described (WEAVER AND DRUMMOND-BARBOSA 2019) by maintaining five experimental females mated with five *AGES* males in perforated plastic bottles capped with molasses/agar plates smeared with 0 mM auxin or the recommended concentration of 10 mM auxin inactive yeast paste. Molasses/agar plates were changed twice daily. The number of eggs laid per day was counted in five replicates per genotype and results were subjected to a paired Student’s *t*-test.

Egg hatching was measured by transferring up to 30 eggs from molasses/agar plates to fresh molasses/agar plates containing 0 mM auxin inactive yeast paste in the center every 2 days and the number of eggs that has hatched were counted 24 hours after the transfer. The number of eggs hatched per day was counted in five replicates per genotype and results were subjected to a paired Student’s *t*-test.

### Adult female ovary immunostaining and fluorescence microscopy

Ovaries and carcasses were dissected in Grace’s Insect Medium (Gibco), fixed, and washed as previously described (WEAVER AND DRUMMOND-BARBOSA 2019). Samples were blocked for at least 3 hours in 5% normal goat serum (NGS; Jackson ImmunoResearch) and 5% bovine serum albumin (BSA; Sigma) in phosphate-buffered saline [PBS; 10 mM NaH_2_PO_4_/NaHPO_4_, 175 mM NaCl (pH 7.4)] containing 0.1% Triton X-100 (PBST). Samples were incubated overnight at 4°C in primary antibodies diluted in blocking solution as follows: mouse monoclonal anti-alpha-spectrin (Developmental Studies Hybridoma Bank; DSHB, 3 µg/ml), mouse monoclonal anti-Lamin C (DSHB, 0.8 µg/ml), and rat monoclonal anti-Vasa (DSHB, 2.15 µg/ml). Samples were washed in PBST and incubated at room temperature for 2 hours with 1:200 Alexa Fluor 488- or 568-conjugated goat-species specific secondary antibodies (Molecular Probes) in blocking solution. Samples were then washed three times for 15 minutes and mounted in Vectashield containing 1.5 µg/ml 4’,6-diamidino-2-phenylindole (DAPI; Vector Laboratories) and imaged using a Leica SP8 confocal.

Cap cells and GSCs were identified as described (WEAVER AND DRUMMOND-BARBOSA 2019), and two-way ANOVA with interaction (GraphPad Prism) was used to calculate the statistical significance of any differences among genotypes in the rate of cap cell or GSC loss from at least three independent experiments, as described (ARMSTRONG *et al*. 2014). Progression through vitellogenesis was assessed using DAPI staining, as described (WEAVER AND DRUMMOND-BARBOSA 2019). Three independent experiments were performed and subjected to a Student’s *t-*test for statistical analysis.

### ApopTag assays

To detect dying germline cysts, the ApopTag Indirect *In Situ* Apoptosis Detection Kit (Millipore Sigma) was used according to the manufacturer’s instructions as previously described (WEAVER AND DRUMMOND-BARBOSA 2019). Briefly, fixed and teased ovaries were rinsed in equilibration buffer twice for 5 minutes each at room temperature. Samples were incubated in 100 µl TdT solution at 37°C for 1 hour with mixing at 15-minute intervals. Ovaries were washed three times in 1X PBS followed by incubation in anti-digoxigenin conjugate for 30 min at room temperature protected from light. Samples were washed four times in 1X PBS and processed for immunofluorescence as described above.

## RESULTS AND DISCUSSION

### Adult *Drosophila* males have increased sensitivity to auxin compared to females

We sought to utilize the AGES expression system in our laboratory for controlling gene expression and began by determining whether we could recapitulate Gal4 expression at levels like that of *Gal80^ts^*. We recombined the auxin-inducible degron line (“*AGES*”) with *3.1Lsp2-Gal4* (*3.1Lsp2-Gal4^AGES^*) to drive the expression of *UAS-nucGFP* in adult female adipocytes compared to the previously described *3.1Lsp2-Gal4* recombined with *Gal80^ts^* (*3.1Lsp2Gal4^ts^*; ARMSTRONG *et al*. 2014). Zero-to-two-day old females of each genotype were fed inactive yeast paste for two days (pre-treatment) prior to Gal4-induction. Gal4 expression was induced in females carrying the *AGES* transgene by feeding solid food supplemented with inactive wet yeast paste containing 0 mM to the recommended dose of 10 mM auxin (inactive yeast was used to prevent auxin metabolism by live yeast) for two days; whereas females carrying the *Gal80^ts^* transgene were shifted from 18°C to 29°C for two days and the relative expression of *GFP* transcripts were measured (**Figure S1A**). There was minimal “leaky” Gal4 expression with *3.1Lsp2-Gal4^AGES^* fed food without auxin (0 mM); whereas 5 mM and 10 mM auxin induced *nucGFP* expression at levels similar to that of *3.1Lsp2-Gal4^ts^*. Furthermore, we found that removal of auxin using *3.1Lsp2-Gal4^AGES^* inactivated Gal4 activity at a faster rate than *3.1Lsp2-Gal4^ts^* (shifted back to 18°C; **Figure S1B**). Therefore, Gal4 expression using AGES can be recapitulated as previously described (MCCLURE *et al*. 2022) and is faster at repressing transgene activation compared to *Gal80^ts^*.

Because the wild-type strain *Canton-S* was previously used to monitor development and survival in flies reared on auxin-containing food (MCCLURE *et al*. 2022), we monitored larval development in flies from the *w^1118^* genotype, a widely-used genetic background strain in *Drosophila* biology. When we measured the time between egg-laying and pupariation in a mixed-sex group of larvae reared on yeast-sugar-cornmeal food supplemented with (5 mM) or without (0 mM) auxin, we found that the time to pupariation was longer in auxin-fed larvae than in larvae raised without auxin (**Figure S1C**). This suggests that the recommended dose of auxin delays larval development in at least one widely-used *Drosophila* strain, indicating that genetic background is an important consideration when using the AGES system for studies at the larval stage of development.

Given these larval phenotypes, we next wanted to determine whether auxin exposure induces physiological changes in adults. Because genetic variation can influence physiology (SHORTER *et al*. 2015; EVANGELOU *et al*. 2019; DAMSCHRODER *et al*. 2020), we used multiple strains to test the effects of auxin on physiology. We first compared the *AGES* line and the strain into which the *AGES* transgene was introduced (*VK00040*) in our experiments. We also used *Oregon-R* [the wild-type strain used for the ModENCODE project (CONSORTIUM *et al*. 2010)] as an additional control. We incubated four-to-six-day old adult males and females of each genotype in vials supplemented with chromatography paper soaked with liquid food containing 0 mM to 350 mM auxin and measured lethality after 48 hours of exposure (**Figure 1A-C; Figure S2A**). Compared to the 0 mM control, both males and females showed increased susceptibility to moderate concentrations of auxin (20-30 mM; **Figure 1B,C; Figure S2A**). In addition, we observed sex differences in sensitivity to auxin exposure, with females able to tolerate higher concentrations relative to males. For example, the benchmark dose (BMD; the concentration that produces a change in the response) in males was lower than that of females in each genotype (**Table 1**). Interestingly, both *AGES* males and females had a higher BMD upon auxin exposure relative to the *VK00040* injection and *Oregon-R* lines, suggesting that the *AGES* transgene may confer resistance to auxin toxicity. These results suggest that exposure to auxin in both males and females results in lethality, where males have a higher susceptibility to auxin exposure compared to females.

**Figure 1.**
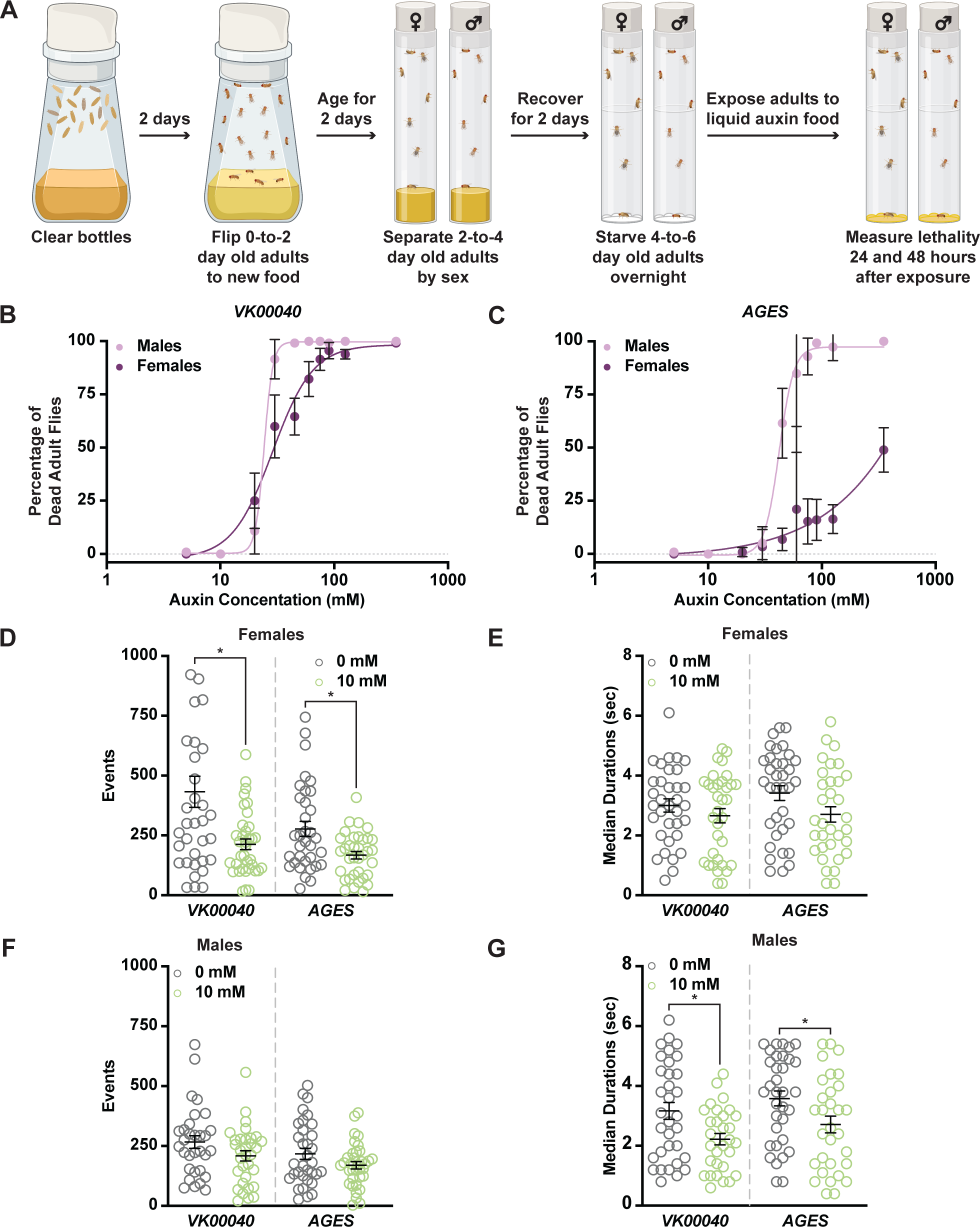
Increased auxin exposure in adult *Drosophila* decreases survival and alters feeding behavior. **(A)** Cartoon schematic of auxin exposure feeding protocol. Cartoon was created using BioRender. **(B,C)** Dose response curves for males and females exposed to increasing concentrations of auxin in the *VK00040* transgene injection line (B) or the *AGES* transgenic line (C). **(D)** The total number of feeding events for females in the *VK00040* or *AGES* fly line exposed to 0 mM or 10 mM auxin. **(E)** The median time of feeding activity of females in the *VK00040* or *AGES* fly lines exposed to 0 mM or 10 mM auxin.**(F)** The total number of events for males in the *VK00040* or *AGES* fly lines exposed to 0 mM or 10 mM auxin. **(G)** The median time of feeding activity of males in the *VK00040* or *AGES* fly lines exposed to 0 mM or 10 mM auxin. Data shown as mean ± SEM. \**P* < 0.05, Mann-Whitney *U*-test.

**Table 1.** Benchmark dose of male and female *Drosophila* exposed to auxin.

| Genotype | Male | Female |
| --- | --- | --- |
| *Oregon-R* | 8.89 mM | 19.97 mM |
| *VK00040* | 17.446 mM | 14.208 mM |
| *AGES* | 32.449 mM | 57.291 mM |

For the remainder of our analyses we used the recommended working concentration of auxin for AGES (10 mM), which as previously described, did not cause lethality in the *AGES* line for either sex [**Figure 1B,C**; **Table 1**; (MCCLURE *et al*. 2022)]. To determine whether feeding behavior could account for sex differences in the survival response to auxin, we analyzed the food consumption of adult male and female flies exposed to 0 mM or 10 mM auxin using the Fly Liquid-Food Interaction Counter [FLIC, (RO *et al*. 2014)] over a 24-hour period (**Figure 1D-G, Figure S2B-E**). In both feeding conditions, females exhibited more feeding events relative to males in all tested genotypes (**Table 2**). For example, females fed 0 mM auxin in both the *VK00040* and *AGES* lines had significantly more engagements with the interaction counter compared to *VK00040* and *AGES* males (compare **Figure 1D** to **Figure 1F**). In addition, genetic background confers differences in feeding behavior as seen by the difference between *VK00040* and *AGES*. Surprisingly, compared to the 0 mM food, females had fewer interactions and slightly decreased event durations when fed the 10 mM auxin food; whereas males only decreased the amount of time they interacted with the food when exposed to 10 mM auxin (**Figure 1G**). These results are consistent with previous reports that females eat more than males on normal food diets with yeast (WONG *et al*. 2009). However, our results also suggest that females exposed to auxin may have an increased survival due to fewer interactions with the food relative to adult males, resulting in differential auxin sensitivity. Therefore, future studies should investigate the female-specific avoidance of auxin, since decreased feeding behavior could confound interpretation of results.

**Table 2.** Average feeding events and durations in male and female *Drosophila*.

| Genotype | Male |  | Female |  |
| --- | --- | --- | --- | --- |
|  | 0 mM | 10 mM | 0 mM | 10 mM |
| *Oregon-R* | 342.8 ± 39.85<br>events<br>3.21 ± 0.26 sec<br>duration | 278.72 ± 26.64<br>events<br>3.21 ± 0.3 sec<br>duration | 519.14 ± 81.52<br>events<br>3.22 ± 0.24 sec<br>duration | 268.82 ± 28.65<br>events<br>3.3 ± 0.29 sec<br>duration |
| *VK00040* | 266.42 ± 26.27<br>events<br>3.17 ± 0.29 sec<br>duration | 208.56 ± 21.15<br>events<br>2.22 ± 0.19 sec<br>duration | 432.23 ± 64.82<br>events<br>3.0 ± 0.22 sec<br>duration | 213.03 ± 22.54<br>events<br>2.66 ± 0.24 sec<br>duration |
| *AGES* | 217.15 ± 23.09<br>events<br>3.58 ± 0.25 sec<br>duration | 169.63 ± 15.25<br>events<br>2.71 ± 0.28 sec<br>duration | 277.44 ± 31.19<br>events<br>3.42 ± 0.25 sec<br>duration | 167.49 ± 15.9<br>events<br>2.71 ± 0.26 sec<br>duration |

### Fatty acid metabolism is decreased in adults exposed to auxin

To determine whether auxin exposure altered metabolism in adult *Drosophila*, we performed metabolomics to compare adult males and females of each genotype exposed to 10 mM auxin relative to the 0 mM control (**Table S1**). Partial Least Squares Discriminant Analysis (PLS-DA) showed that each genotype clustered in distinct groups (**Figure S3**). Notably, 0 mM controls clustered separately from 10 mM auxin samples in all genotypes for both sexes along the Component 1 axis, which describes 33.4% or 34.4% of the variance for females and males, respectively. We further analyzed the data using MetaboAnalyst (PANG *et al*. 2021), which revealed that the levels of acylcarnitine and fatty acid metabolites were the most significantly decreased metabolites in females exposed to 10 mM auxin regardless of genotype (**Figure 2A; Figures S4-6**). Adult males exposed to 10 mM auxin also exhibited decreased fatty acid and acylcarnitine metabolites (**Figure 2B**); however, amino acid levels were the most significantly altered metabolites in males in response to auxin (**Figures S4-6**).

**Figure 2.**
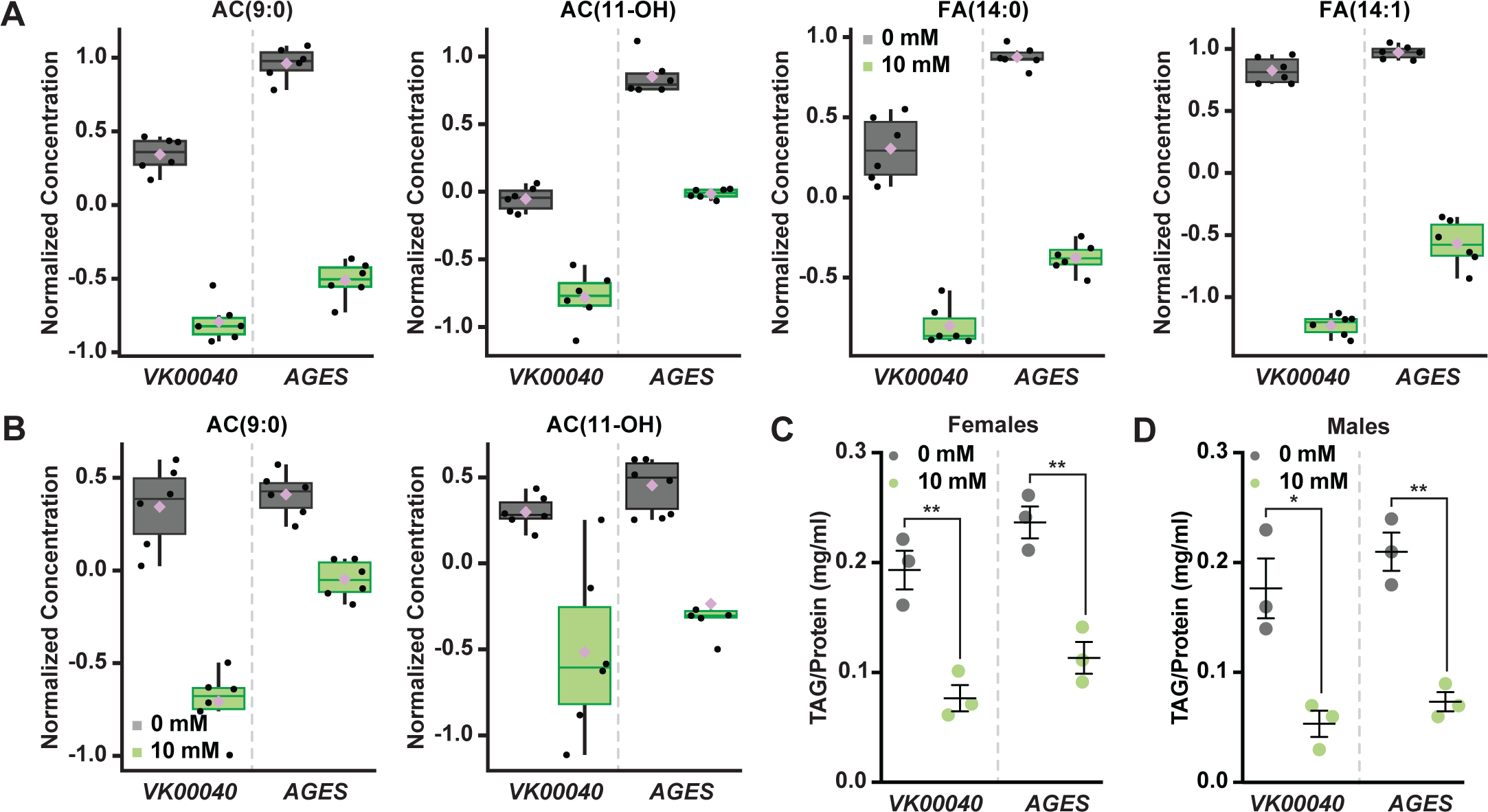
Auxin exposure decreases acylcarnitine, fatty acid, and triglyceride levels in adult *Drosophila*. **(A,B)** Box plots illustrating the relative abundance of acylcarnitine or fatty acid metabolites in adult females (A) or males (B) exposed to 0 mM or 10 mM auxin. All box plots were generated using MetaboAnalyst 5.0 as described in the methods. Black dots represent individual samples, the horizontal bar in the middle represents the median, and the purple diamond represents the mean concentration. For all box plots, the metabolite fold-change was >2-fold and *P* < 0.05. **(C,D)** TAG contents (mg TAG/mg protein) in whole adult females (C) or males (D) in the *VK00040* or *AGES* fly lines exposed to 0 mM or 10 mM auxin. Data shown as mean ± SEM. \**P* < 0.05, \*\**P* < 0.01, two-tailed Student’s *t*-test.

To confirm that fatty acid metabolism is altered in response to auxin, we exposed adult males and females of each genotype to 0 mM or 10 mM auxin and measured the levels of triacylglycerol (TAG) after 48 hours of exposure. Both males and females exposed to 10 mM auxin of each genotype had significantly less TAG relative to the 0 mM control (**Figure 2C,D; Figure S4G**). We note that the lack of sex difference in lipid levels is likely due to exposing adults to auxin prior to the onset of male-female differences in TAG (WAT *et al*. 2020). To determine whether auxin influences TAG in additional contexts, we transferred newly-eclosed virgin *w^1118^* males and females to a yeast-sugar-cornmeal diet supplemented with either 0 mM or the recommended dose of 10 mM auxin and measured TAG levels after 5 days. We found a significant decrease in whole-body fat storage in both male and female flies exposed to 10 mM auxin compared with flies transferred to food with 0 mM auxin (**Figure S4H**); however, the magnitude of the auxin-induced decrease in body fat was greater in females than in males (sex:diet interaction *p* = 0.001; two-way ANOVA). This suggests that auxin has a stronger effect on female TAG levels, in line with our data showing a female-biased reduction in food interactions on auxin-supplemented medium.

We next asked whether this effect on TAG levels was reversible. We exposed 0-day-old male and female flies to diets with 0 mM or 10 mM auxin for five days, and then measured TAG levels after shifting the flies to food supplemented with no auxin for five additional days. We found that females but not males showed a strong trend toward recovery of whole-body TAG levels after shifting the flies back to food with no auxin (Figure S4I), suggesting the effect of auxin feeding on body fat were reversible only in females. Furthermore, we found that flies exposed to food supplemented with 10 mM auxin for 10 days show no additional reduction in whole-body TAG levels compared with flies exposed to auxin for 5 days in either sex (Figure S4I). Taken together, these results suggest that auxin exposure disrupts fatty acid metabolism, resulting in decreased circulating lipids in adult males and females. While these changes to fat metabolism are partially reversible after withdrawal of auxin supplementation in females, changes to whole-body TAG levels persisted even after auxin withdrawal in males.

### Auxin exposure induces global transcriptomic changes in adult *Drosophila*

To determine whether auxin exposure induced changes in gene expression, we exposed adult males and females of each genotype to 0 mM and 10 mM auxin for 48 hours and performed RNA sequencing analysis of whole animals. In both males and females of each tested genotype, at least 150 transcripts were significantly altered in response to auxin exposure (**Figures 3A-D and S7-8; Tables S2-7**). Genes involved in drug metabolism [e.g., glutathione-S-transferases (GSTs) and uridine diphosphate-glucuronosyltransferases (UGTs), phase II enzymes required to increase hydrophobicity of compounds (YU 2008)] were significantly up-regulated in response to auxin exposure in both males and females of all genotypes (**Figure 3E; Figures S7-8**), suggesting that auxin induces a xenobiotic response at this concentration (YU 2008). Surprisingly, genes involved in fatty acid metabolism (e.g., Fad2, which encodes a desaturase) were also significantly up-regulated; whereas lipases (enzymes that break down fatty acids) were significantly down-regulated in each genotype and sex (**Figure 3F; Figure S7D; Figure S8C,D**). Notably, the transcription of enzymes associated with lipolysis such as the lipase *brummer* (GRONKE *et al*. 2005) or the perilipins *Lsd-1* and *Lsd-2* (BELLER *et al*. 2010) were unaltered in response to auxin exposure (**Tables S2-7**). Therefore, it is possible that in response to auxin, triglycerides are mobilized by an unknown mechanism resulting in upregulation of fatty acid metabolism and attenuation of lipid breakdown. It is also possible that decreased interactions or feeding time with auxin-containing food promotes fasting, resulting in decreased TAG and lipid synthesis. Collectively, these results suggest that auxin exposure in adults significantly alters the transcriptome to adjust to decreased fatty acid metabolism and lipid stores.

**Figure 3.**
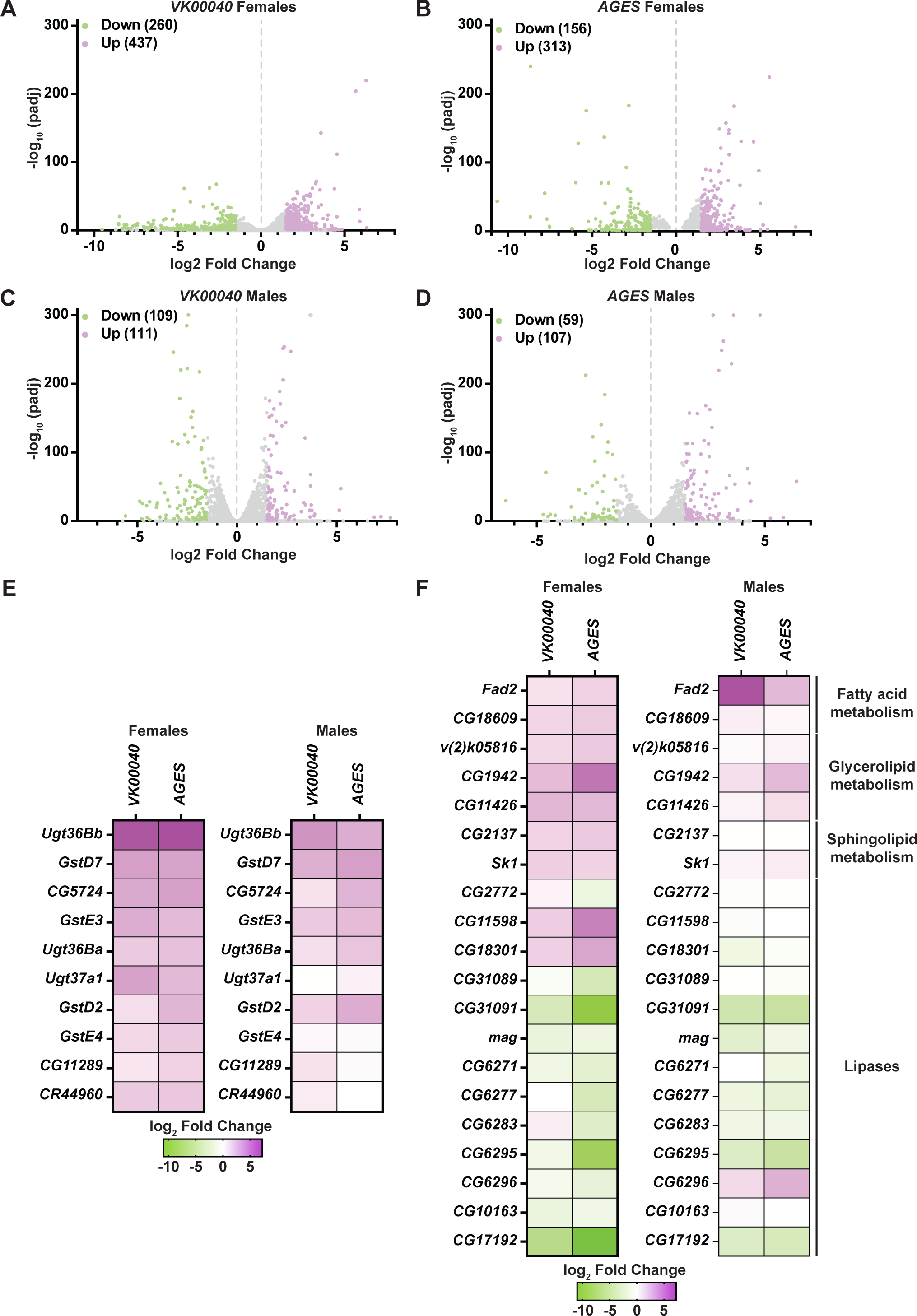
Lipid metabolism and detoxification transcripts are significantly altered in adult *Drosophila* in response to auxin exposure. **(A-D)** Volcano plots of differentially expressed genes graphing the statistical significance [-log_10_(padj)] against the magnitude of differential expression (log_2_ Fold Change) in females (A,B) or males (C,D) of the *VK00040* or *AGES* line. **(E)** Significantly up-regulated genes involved in “drug metabolism” in adult females or males of the *VK00040* or *AGES* lines exposed to 10 mM auxin and compared to the 0 mM control. **(F)** Significantly up-(purple) and down-regulated (green) genes involved in fatty acid metabolism in adult females or males of the *VK00040* or *AGES* lines exposed to 10 mM auxin and compared to the 0 mM control.

Auxin is an essential hormone for plant growth and development (DU *et al*. 2020; WOJCIK *et al*. 2020; GOMES AND SCORTECCI 2021), and has been widely used to manipulate gene expression in multiple organisms (ZHANG *et al*. 2015; TROST *et al*. 2016; CHEN *et al*. 2018; LI *et al*. 2019; SHETTY *et al*. 2019; YESBOLATOVA *et al*. 2020; MACDONALD *et al*. 2022). In *Arabidopsis*, auxin is a known regulator of fatty acid synthesis in plants (HE *et al*. 2020) and has been shown to induce lipid synthesis to control vacuole trafficking (LI *et al*. 2015). Our results suggest that auxin has the opposite effect on lipid metabolism in adult *Drosophila* and decreases triglyceride levels. Given its widespread use, the role of auxin in potentially regulating lipid metabolism should be further characterized in additional model organisms using auxin to manipulate gene expression (at their respective recommended dosages), such as *C. elegans* (ZHANG *et al*. 2015) and mice (MACDONALD *et al*. 2022).

### Auxin exposure in adult females does not influence oogenesis

Although many tissues maintain and regulate lipid stores in *Drosophila* (including muscle and gut [reviewed in (HEIER AND KUHNLEIN 2018)]), the adult adipose tissue is a major lipid storage depot [reviewed in (CHATTERJEE AND PERRIMON 2021)] and impacts peripheral tissue function such as adult *Drosophila* oogenesis (ARMSTRONG *et al*. 2014; MATSUOKA *et al*. 2017; ARMSTRONG AND DRUMMOND-BARBOSA 2018; WEAVER AND DRUMMOND-BARBOSA 2018; WEAVER AND DRUMMOND-BARBOSA 2019). Thus, we sought to examine if the metabolic effects of auxin described above impact oogenesis.

Each ovary consists of 16-20 ovarioles composed of progressively older follicles that ultimately give rise to a mature egg chamber [**Figure 4A,B**; (DRUMMOND-BARBOSA 2019)]. Oogenesis is maintained by 2-3 germline stem cells (GSCs) that reside in the anterior germarium of each ovariole and can be identified based on their proximity to the stem cell niche, which is primarily composed of somatic cap cells. GSCs self-renew and give rise to early GSC progeny that differentiate to produce follicles that bud from the germarium and complete oogenesis.

**Figure 4.**
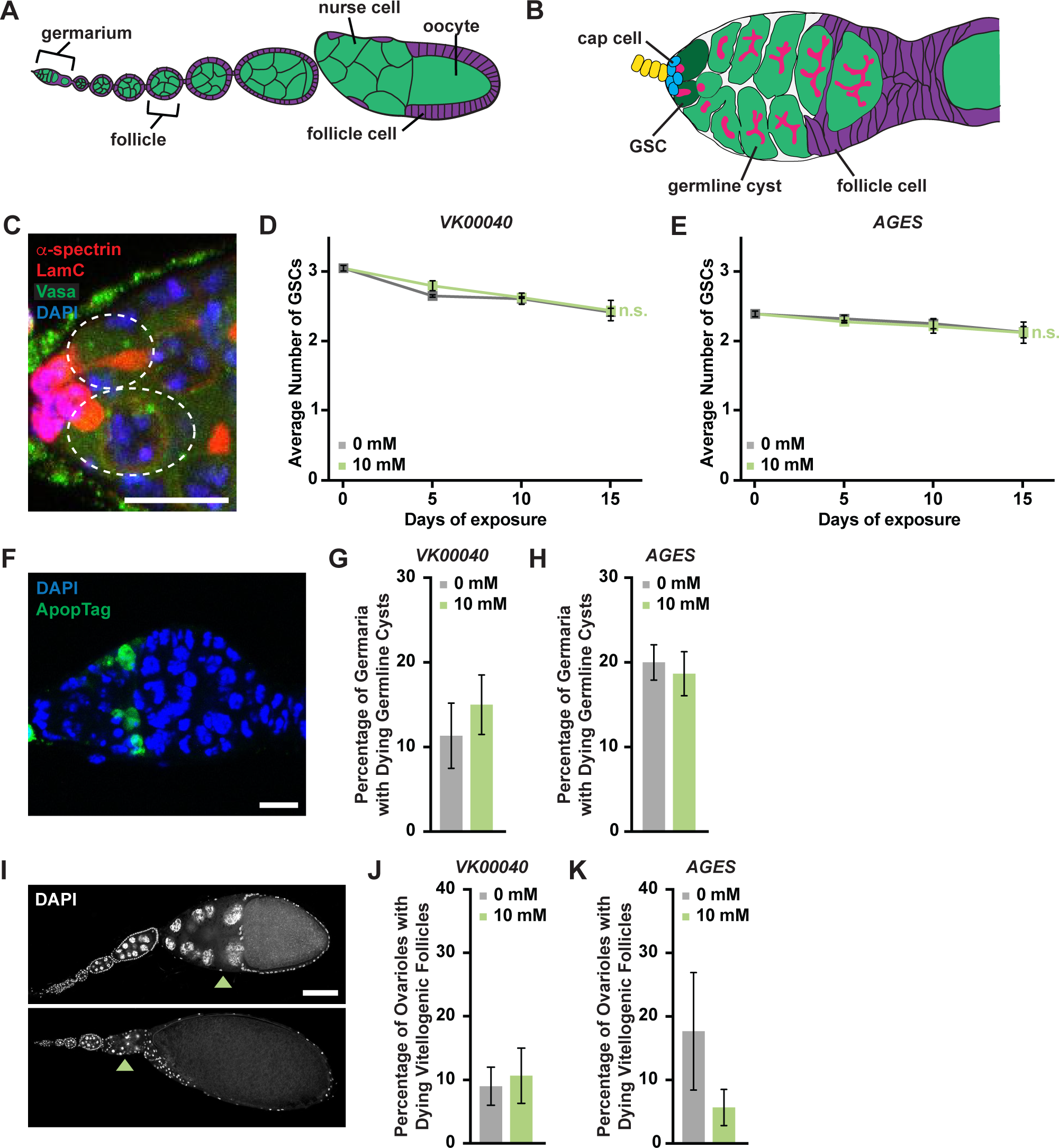
Auxin exposure does not significantly influence processes of oogenesis. **(A)** Cartoon schematic of the adult *Drosophila* ovariole showing the anterior germarium followed by developing egg chambers, which consist of 16 germ cells (15 nurse cells and one oocyte; green) that are surrounded by follicle cells (purple). **(B)** Schematic of the germarium, which contains 2-3 germline stem cells (GSCs, dark green) and somatic cells (gray and purple). Each GSC divides asymmetrically to self-renew and generate a cystoblast that divides to form a 16-cell cyst. Early germline cysts are surrounded by follicle cells (purple) to bud a new egg chamber. GSCs and their progeny are identified based on the position and morphology of the fusome (pink), a germline-specific organelle. **(C)** Germaria from adult females exposed to 10 mM auxin for 10 days. α-spectrin (magenta), fusome; LamC (red), cap cell nuclear lamina; Vasa (green), germ cells; DAPI (blue), nuclei. GSCs are outlined. Scale bar, 10 µm. **(D,E)** Average number of GSCs per germarium over time in females exposed to 0 mM or 10 mM auxin in the *VK00040* (D) *AGES* (E) lines. Data shown as mean ± SEM. No statistically significant differences, two-way ANOVA with interaction. **(F)** Germaria from adult females exposed to 10 mM auxin. ApopTag (green), dying cells; DAPI (blue), nuclei). **(G,H)** Average percentage of germaria containing ApopTag-positive germline cysts in adult females in the *VK00040* (G) or *AGES* (H) lines exposed to 0 mM or 10 mM auxin. Data shown as mean ± SEM; Student’s *t*-test. 100 germaria were analyzed for each genotype and condition. **(I)** Ovarioles exposed to 10 mM auxin for 10 days showing a healthy ovariole (top) and an ovariole with a dying vitellogenic egg chamber (bottom). Arrowheads point to healthy or dying vitellogenic egg chambers. DAPI (white), nuclei. Scale bar, 100 µm. **(J,K)** Average percentages of ovarioles containing dying vitellogenic egg chambers in females exposed to 0 mM or 10 mM auxin in the *VK00040* (J) or *AGES* (K) lines. Data shown as mean ± SEM, Student’s *t*-test. 100 ovarioles were analyzed for each genotype and condition.

To determine if auxin exposure influences fecundity or progeny survival, we performed egg laying and larval hatching analyses. Adult females (*Oregon-R*, *VK00040*, or *AGES*) were maintained with *AGES* males (due to their increased resistance to auxin) on molasses plates supplemented with wet inactive yeast containing either 0 mM auxin or 10 mM auxin for 15 days. Relative to 0 mM auxin control, exposure to 10 mM auxin did not significantly influence the number of eggs laid per female in any tested genotype (**Figure S9A,C,E**). We note that the number of eggs laid under 0 mM auxin conditions is relatively low; however, this is likely due to the decrease in egg production in females fed inactive yeast compared to active yeast paste (**Figure S10**). In addition, the relative hatching percentages of each genotype also were not affected by auxin exposure (**Figure S9B,D,F**). These results suggest that auxin exposure in adult females does not significantly influence the number of laid eggs or oocyte quality.

We next analyzed specific processes of oogenesis that are sensitive to metabolic changes including GSC maintenance, early germline cyst survival, and survival of vitellogenic follicles (DRUMMOND-BARBOSA 2019). We detected GSCs by the morphology and location of the fusome relative to the GSC-niche (**Figure 4C**). GSC numbers in all genotypes exposed to 10 mM auxin were comparable to those in 0 mM auxin controls at each time point (**Figure 4D-E; Figure S11A,D**). Likewise, there was no change in the number of cap cells over time (**Figure S11E**). Furthermore, there were no effects of auxin exposure on the survival of early germline cysts [as determined by ApopTag TUNEL labeling (Drummond-Barbosa and Spradling, 2001)] (**Figure 4F-H; Figure S11B**) or survival of vitellogenic egg chambers (**Figure 4I-K; Figure S11C**). Based on these data, we conclude that the recommended auxin concentration for AGES in adult females does not adversely affect oogenesis, despite significant decreases in whole-body fatty acid metabolism and lipid composition.

Consistent with our results, it was recently shown that obese adult *Drosophila* females do not have reduced fecundity, but that fertility defects manifest only when combined with a high sugar diet (NUNES AND DRUMMOND-BARBOSA 2023). Collectively, these results suggest that lipid content alone (lean or obese) is not sufficient to regulate distinct processes of oogenesis in adult *Drosophila* females. Therefore, AGES may be suitable for the study of some processes in adult oogenesis.

### Conclusions and suggestions for future studies

The *Gal4/UAS* system has revolutionized the ability to perform tissue specific manipulations in *Drosophila*. However, finding the ideal conditions to manipulate tissues temporally without causing significant alterations in physiology or impacting organism behavior presents a challenge. For example, using *Gal4*/*UAS* in conjunction with *Gal80^ts^* for adult specific manipulations has the adverse effect of decreasing adult female fecundity, making this system unsuitable for aging studies on oogenesis (GANDARA AND DRUMMOND-BARBOSA 2022). Similarly, controlling transgene expression using RU486 inducible *GeneSwitch* drivers has not only been found to be less than ideal for feeding and aging studies (LANDIS *et al*. 2015; YAMADA *et al*. 2017), but also presents a workplace hazard to pregnant researchers (AVRECH *et al*. 1991). Likewise, our findings herein indicate that methods requiring auxin for temporal transgene induction or protein degradation may work well to study many biological processes, but may not be the ideal system for the study of lipogenesis or other metabolic processes dependent on lipid content. Indeed, our data suggests that auxin-induced changes to metabolism persist even after its withdrawal from the diet, indicating that even short-term auxin treatments may have unwanted physiological effects.

Each modification to the *Gal4/UAS* system designed to provide temporal regulation of transgene expression has strengths and weaknesses and should be carefully vetted to ensure the most ideal experimental design is balanced with the caveats of the methodology. Regardless of the system used, researchers should determine whether their experimental manipulation alters Gal4 expression patterns compared to control conditions [i.e., thoroughly analyze Gal4 expression pattern across developmental time and tissues (WEAVER *et al*. 2020)]. For example, the expression of transgenes using numerous neuronal Gal4 drivers with AGES is significantly weaker than Gal4-induced expression at 30°C (HAWLEY *et al*. 2023). Finally, researchers should ensure that non-specific transgene controls used in the same conditions as the experimental (e.g., 0 mM versus 10 mM auxin) do not have phenotypes for physiological outputs of interest and ensure that changes in physiology that could confound result interpretations are accounted for.

## DATA AVAILABILITY

*Drosophila* strains can be purchased from the Bloomington *Drosophila* Stock Center. The data and analyses in this paper are described in the main figures. The raw data and processed data files are available through the NCBI GEO accession number GSE237283 and are also provided as supplemental figures and tables. Additional raw data is available upon request.

## ACKNOWLEDGEMENTS

We thank the Bloomington Stock Center (National Institutes of Health P40OD018537) for *Drosophila* stocks. We are thankful to Flybase (www.flybase.org), an essential *Drosophila* research resource (NIH 5U41HG000739). The authors would like to thank the Indiana University Light Microscopy Imaging Center (LMIC) for access to microscopy facilities. We are grateful for Kristina J. Weaver and Scott D. Pletcher for assistance with the FLIC code. We are also grateful to Elizabeth T. Ables, Brian R. Calvi, and Deepika Vasudevan for critical reading of the manuscript. This work was supported by the Canadian Institutes for Health Research grant PJT-153072 (E.J.R. and P.B.), the National Institutes of Health (NIH) grants R35 GM119557 (J.M.T), R00 GM127605 (L.N.W.), and R35 GM150517 (L.N.W).

## FUNDING

This work was supported by the Canadian Institutes for Health Research grant PJT-153072 (E.J.R and PB), the National Institutes of Health (NIH) grants R35 GM119557 (J.M.T), R00 GM127605 (L.N.W.), and R35 GM150517 (L.N.W).

## CONFLICT OF INTEREST

The authors declare no competing interests.

## AUTHOR CONTRIBUTIONS

S.A.F., P.B., E.D.D., R.L.K., R.C.E., T.N., A.D’A., and L.N.W. performed experiments, analyzed, and interpreted the data. L.N.W., P.B., and E.J.R. wrote the manuscript and J.M.T provided edits.

## Notes

### Competing Interest Statement

The authors have declared no competing interest.

### Summary of Updates

The main text has been updated to include new results. There is a new supplemental figure 8 and an additional subpanel to supplemental figure 4. In addition, author emails and ORCID numbers have been altered.

