## Supplemental Figures for "Auxin Exposure Disrupts Feeding Behavior and Fatty Acid Metabolism in Adult *Drosophila*"

**This file includes**:

Figures S1 to S11

### SUPPLEMENTARY FIGURES

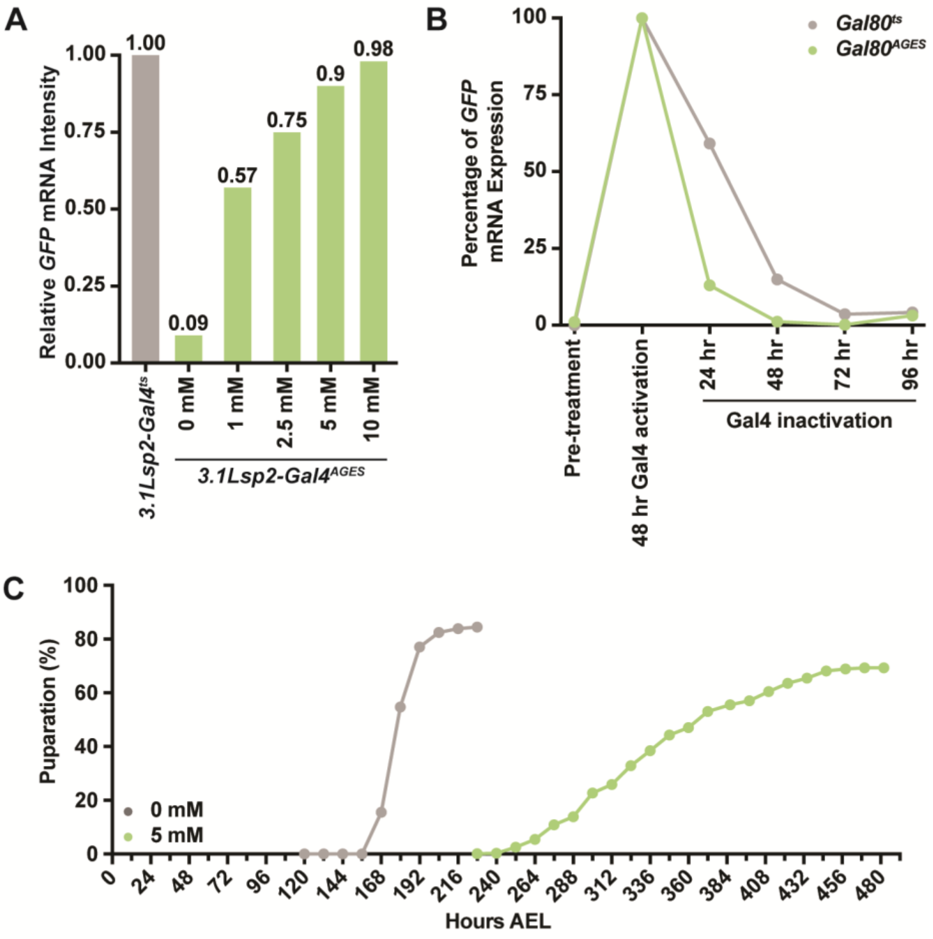


**Figure S1. Auxin exposure increases the time of development.**

**(A)** Relative mRNA intensity of *GFP* transcripts from *3.1Lsp2-Gal4*$^{ts}$ or *3.1Lsp2-Gal4*$^{AGES}$ to induce Gal4-mediated transgene expression. **(B)** Relative *GFP* transcript amount from *3.1Lsp2-Gal4*$^{ts}$ or *3.1Lsp2-Gal4*$^{AGES}$ during Gal4 activation and suppression. **(C)** Time to pupariation indicated in mixed-sex group of *w*$^{1118}$ larvae reared on yeast-sugar-cornmeal food supplemented with 0 mM or 5 mM auxin. Time to pupariation was longer in *w*$^{1118}$ auxin-exposed larvae than in larvae reared on a diet without auxin.

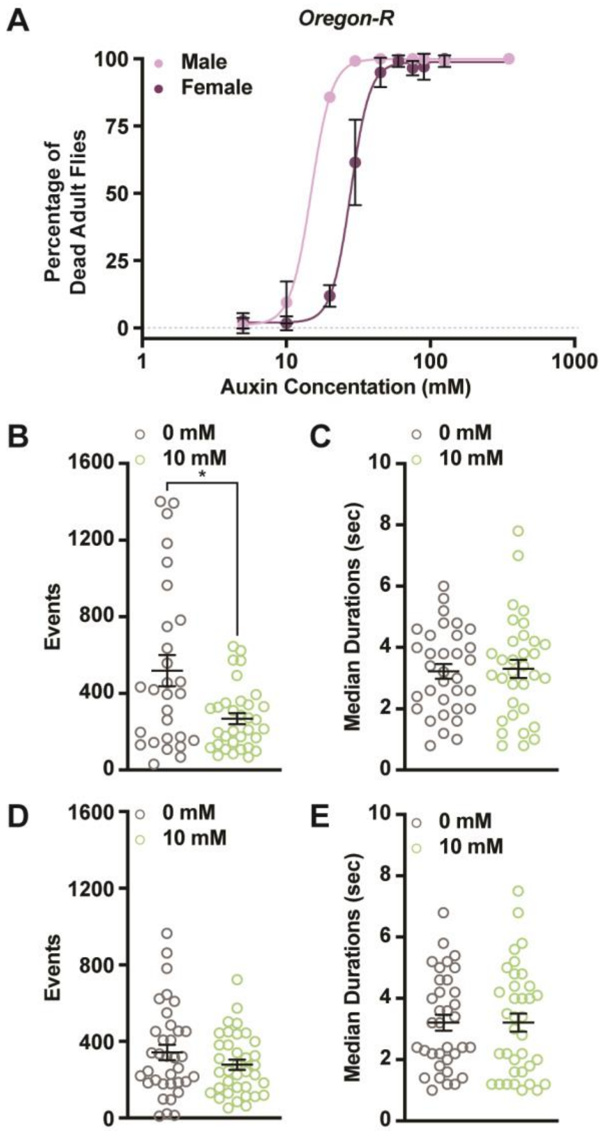


**Figure S2. Auxin exposure in *Oregon-R* adults increases lethality and alters female feeding behavior.**

**(A)** Lethality curves for adult *Oregon-R* males and females exposed to increasing concentrations of auxin. **(B)** The total number of feeding events in *Oregon-R* females exposed to 0 mM or 10 mM auxin. **(C)** The median time of feeding activity in *Oregon-R* females exposed to 0 mM or 5 mM auxin. **(D)** The total number of events in *Oregon-R* males exposed to 0 mM or 10 mM auxin. **(E)** The median time of feeding activity in

*Oregon-R* males exposed to 0 mM or 10 mM auxin. Data shown as mean ± SEM. *$P <$ 0.05, Mann-Whitney *U*-test.

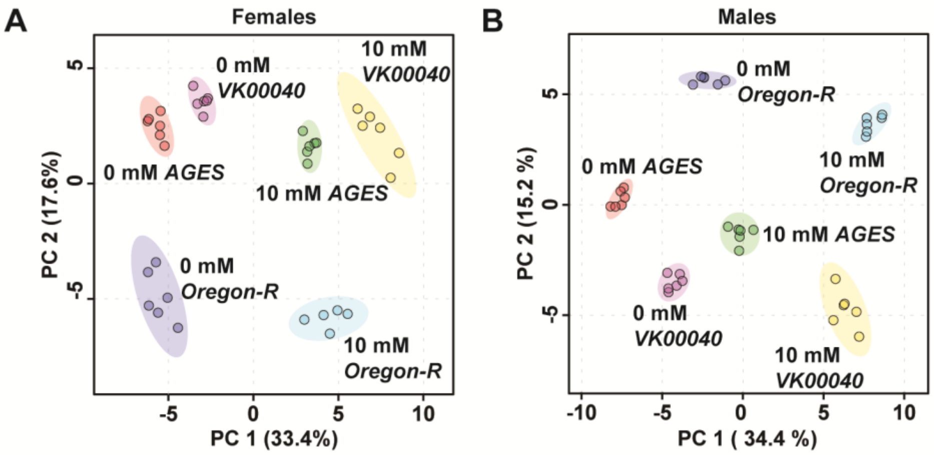


**Figure S3. Auxin exposure alters metabolism in adult *Drosophila* males and females.**

**(A,B)** Comparison of the metabolomic profile from adult females (A) or males (B) exposed to 0 mM or 10 mM auxin in the *Oregon-R*, *VK00040*, or *AGES* background. Metabolomics data from Table S1 was analyzed using principal component analysis conducted by MetaboAnalyst 5.0.

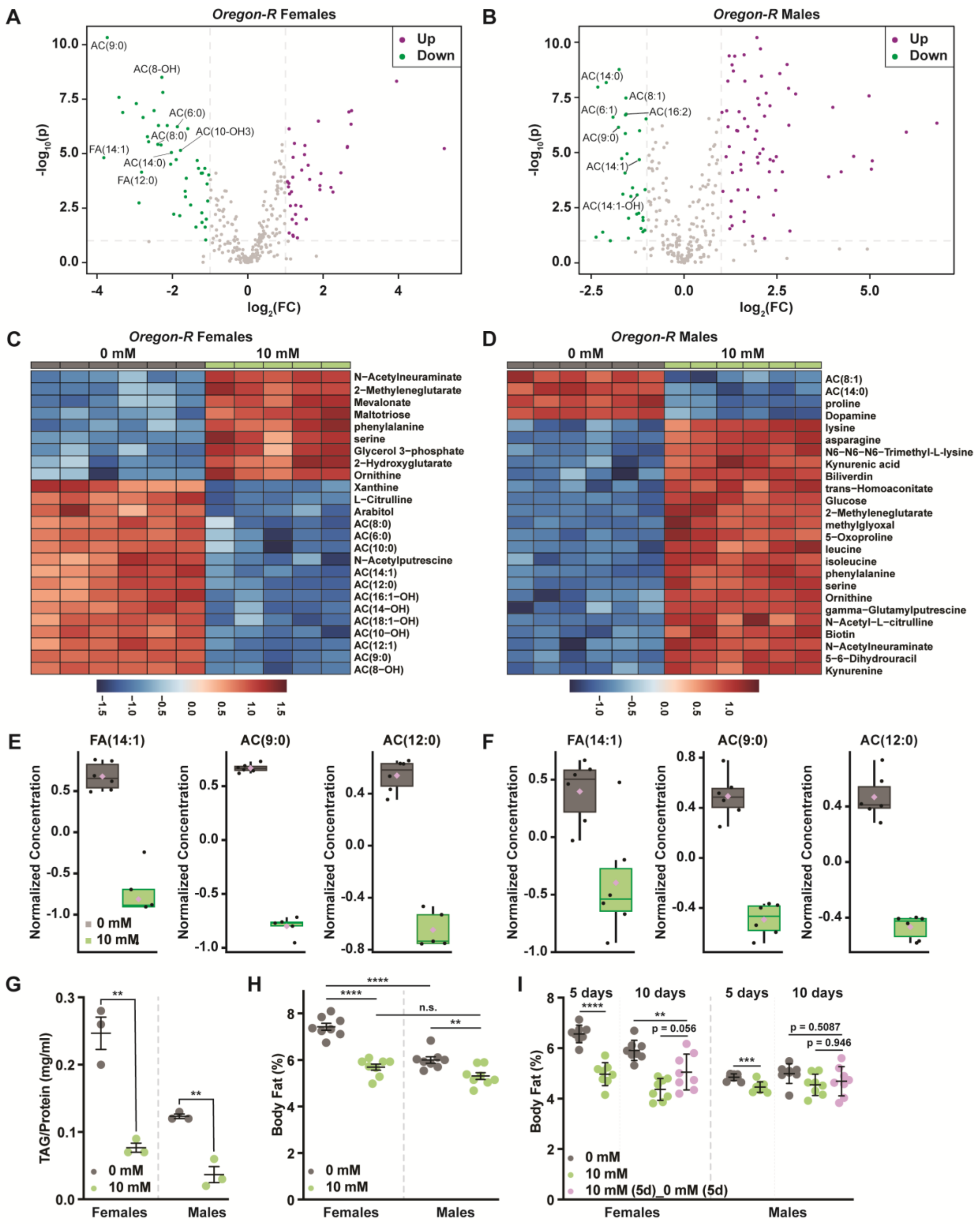

A
Oregon-R Females
Up
Down
AC(9:0)
AC(8-OH)
AC(6:0)
AC(10-OH3)
AC(8:0)
FA(14:1)
AC(14:0)
FA(12:0)
-log10(p)
log2(FC)
B
Oregon-R Males
Up
Down
AC(14:0)
AC(8:1)
AC(6:1)
AC(16:2)
AC(9:0)
AC(14:1)
AC(14:1-OH)
-log10(p)
log2(FC)
C
Oregon-R Females
0 mM
10 mM
N−Acetylneuraminate
2−Methyleneglutarate
Mevalonate
Maltotriose
phenylalanine
serine
Glycerol 3−phosphate
2−Hydroxyglutarate
Ornithine
Xanthine
L−Citrulline
Arabitol
AC(8:0)
AC(6:0)
AC(10:0)
N−Acetylputrescine
AC(14:1)
AC(12:0)
AC(16:1−OH)
AC(14−OH)
AC(18:1−OH)
AC(10−OH)
AC(12:1)
AC(9:0)
AC(8−OH)
D
Oregon-R Males
0 mM
10 mM
AC(8:1)
AC(14:0)
proline
Dopamine
lysine
asparagine
N6−N6−N6−Trimethyl−L−lysine
Kynurenic acid
Biliverdin
trans−Homoaconitate
Glucose
2−Methyleneglutarate
methylglyoxal
5−Oxoproline
leucine
isoleucine
phenylalanine
serine
Ornithine
gamma−Glutamylputrescine
N−Acetyl−L−citrulline
Biotin
N−Acetylneuraminate
5−6−Dihydrouracil
Kynurenine
E
FA(14:1)
AC(9:0)
AC(12:0)
Normalized Concentration
0 mM
10 mM
F
FA(14:1)
AC(9:0)
AC(12:0)
Normalized Concentration
G
TAG/Protein (mg/ml)
0 mM
10 mM
Females
Males
H
Body Fat (%)
0 mM
10 mM
n.s.
Females
Males
I
5 days
10 days
5 days
10 days
p = 0.056
p = 0.5087
p = 0.946
Body Fat (%)
0 mM
10 mM
10 mM (5d)_0 mM (5d)
Females
Males

**Figure S4. Acylcarnitine and fatty acid levels decrease in *Oregon-R* adult males and females exposed to auxin.**

**(A,B)** Volcano plot showing the differences in metabolite abundance in *Oregon-R* females (A) or *Oregon-R* males (B) exposed to 10 mM auxin and compared to the 0 mM auxin control as determined from MetaboAnalyst 5.0 analysis. Dashed vertical lines represent a $\log_2$ fold change (FC) greater than 1.5. The horizontal dashed line represents $P < 0.01$. **(C,D)** Heat maps illustrating the top 25 most significantly altered compounds in *Oregon-R* females (C) or *Oregon-R* males (D) exposed to 10 mM auxin compared to 0 mM auxin controls as determined from MetaboAnalyst 5.0 analysis. **(E,F)** Box plots illustrating the relative abundance of acylcarnitine or fatty acid metabolites in adult *Oregon-R* females (E) or males (F) exposed to 0 mM or 10 mM auxin. Black dots represent individual samples, the horizontal bar in the middle represents the median, and the purple diamond represents the mean concentration. For all box plots, plots were generated using MetaboAnalyst 5.0 as described in the methods, the metabolite fold-change was $>2$-fold, and $P < 0.01$. **(G)** TAG contents (mg TAG/mg protein) in whole adult *Oregon-R* females or males exposed to 0 mM or 10 mM auxin. Data shown as mean ± SEM. $^{**}P < 0.01$, two-tailed Student's *t*-test. **(H)** Percent body fat in virgin 5-day-old $w^{1118}$ male and female flies. Larvae were reared on sugar-yeast-cornmeal food and transferred to food supplemented with 0 mM or 10 mM auxin at eclosion. Both females and males showed a significant decrease in percent body fat when maintained on a diet supplemented with the recommended auxin concentration ($P < 0.0001$ and $P = 0.01$, respectively; two-way ANOVA). Adult females showed a greater reduction in body fat compared with males in response to auxin exposure (sex:diet interaction $P = 0.001$;

two-way ANOVA). (I) Percent body fat in virgin $w^{1118}$ male and female flies. Larvae were reared on sugar-yeast-cornmeal food, and transferred to three different diets at eclosion: (1) sugar-yeast-cornmeal diet with no auxin, (2) sugar-yeast-cornmeal diet supplemented with 10 mM auxin for 5 days then shifted to sugar-yeast-cornmeal food with 0 mM auxin for 5 days, (3) sugar-yeast-cornmeal diet supplemented with 10 mM auxin for 10 days. Flies were collected for analysis at 5 days and 10 days post-eclosion. Adult females showed a greater reduction in body fat compared with males in response to auxin exposure (sex:diet interaction $P < 0.0001$; two-way ANOVA). While percent body fat showed a strong trend toward recovery from 5 days of auxin treatment in females ($P = 0.056$; one-way ANOVA), body fat was still lower than in control females that had not been exposed to auxin ($P = 0.0095$, one-way ANOVA). This trend toward recovery in body fat was not observed in males ($P = 0.946$, one-way ANOVA). Prolonged (10 day) exposure to 10 mM auxin caused no further decrease in body fat compared with a shorter (5 day) exposure in either females or males ($P = 0.1156$ and 0.9890, respectively; one-way ANOVA). **$P < 0.01$, ***$P < 0.01$, ****$P < 0.0001$, two-tailed Student's *t*-test.

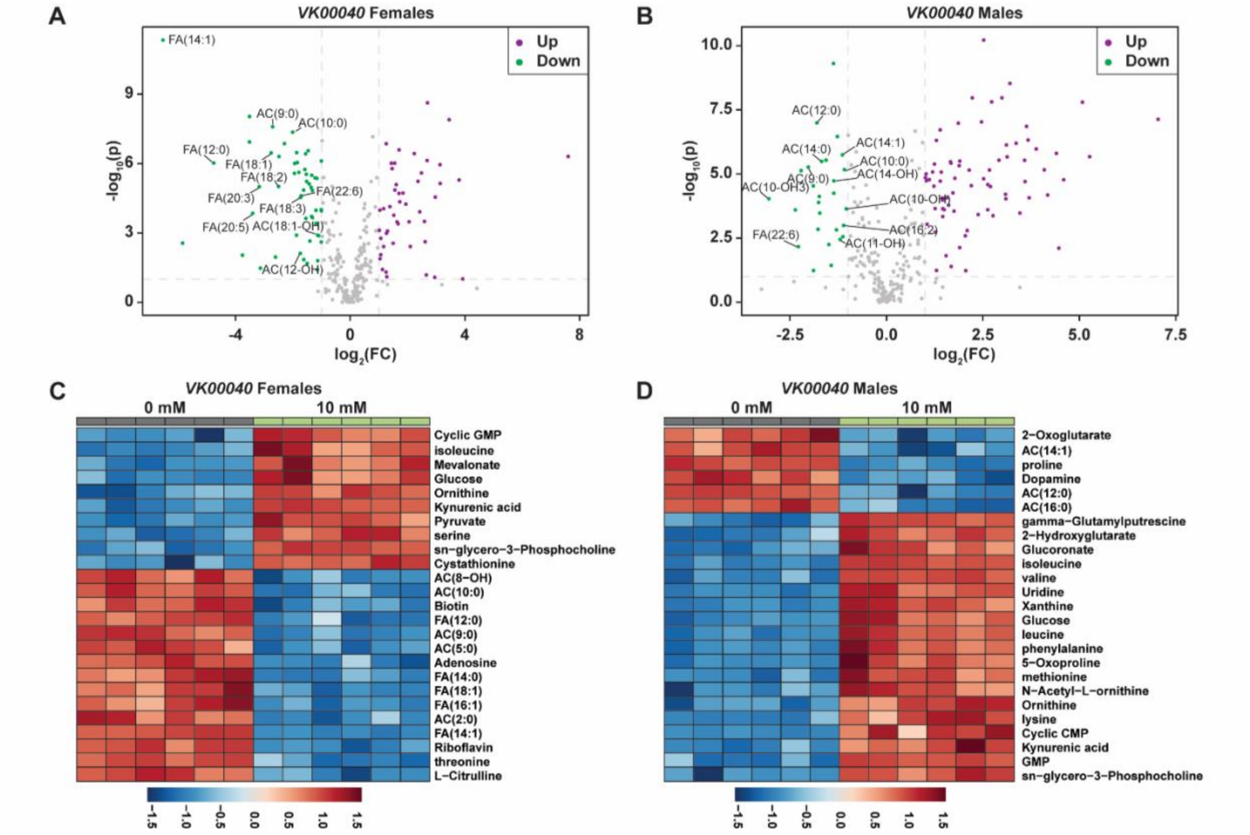


**Figure S5. Auxin exposure decreases acylcarnitine and fatty acid metabolites in *VK00040* adult males and females exposed to auxin.**

**(A,B)** Volcano plot showing the differences in metabolite abundance in *VK00040* females (A) or *VK00040* males (B) exposed to 10 mM auxin and compared to the 0 mM auxin control as determined from MetaboAnalyst 5.0 analysis. Dashed vertical lines represent a $\log_2$ fold change (FC) greater than 1.5. The horizontal dashed line represents $P < 0.01$. **(C,D)** Heat maps illustrating the top 25 most significantly altered compounds in *VK00040* females (C) or *VK00040* males (D) exposed to 10 mM auxin compared to 0 mM auxin controls as determined from MetaboAnalyst 5.0 analysis.

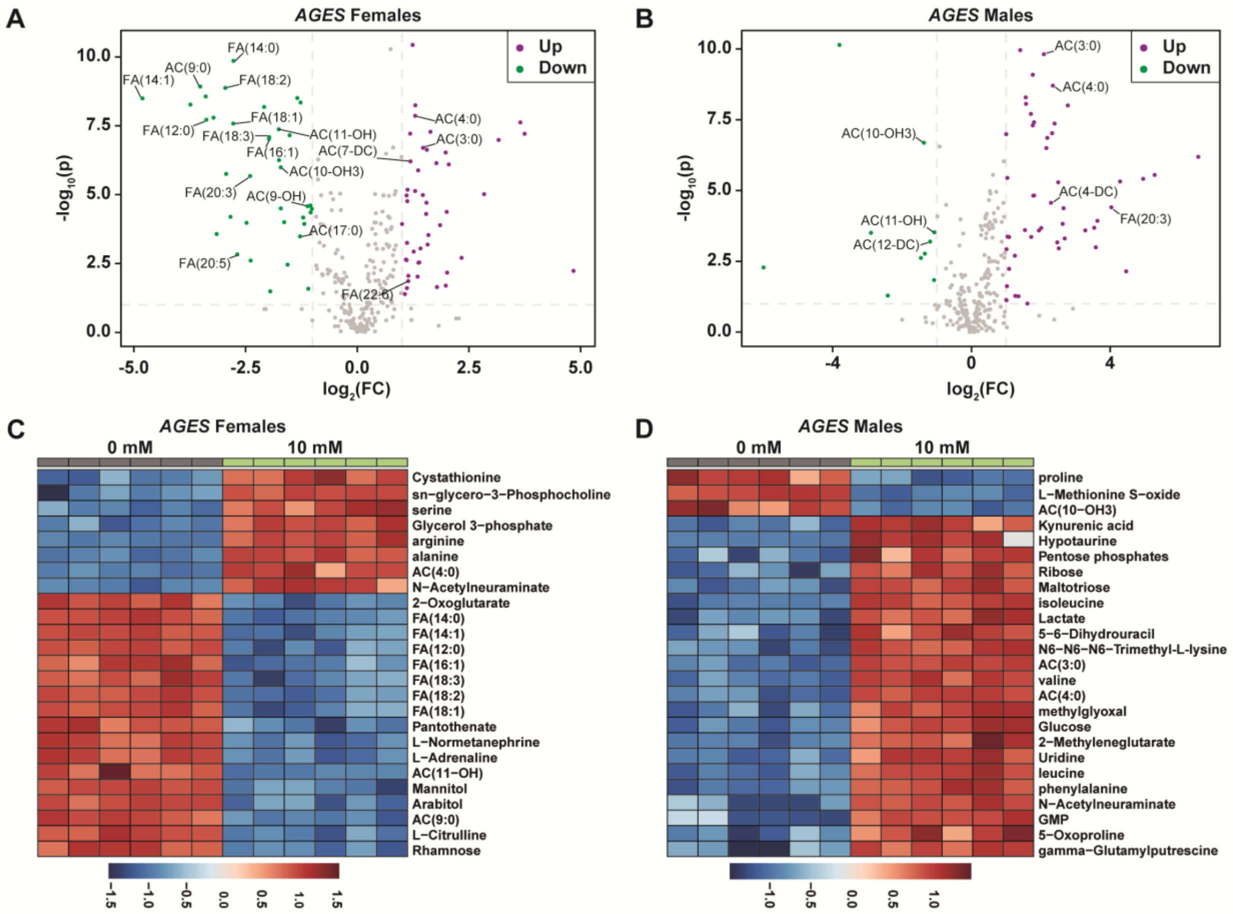


**Figure S6. Acylcarnitine and fatty acid levels decrease in *AGES* adult males and females exposed to auxin.**

**(A,B)** Volcano plot showing the differences in metabolite abundance in *AGES* females (A) or *AGES* males (B) exposed to 10 mM auxin and compared to the 0 mM auxin control as determined from MetaboAnalyst 5.0 analysis. Dashed vertical lines represent a $\log_2$ fold change (FC) greater than 1.5. The horizontal dashed line represents $P <$ 0.01. **(C,D)** Heat maps illustrating the top 25 most significantly altered compounds in *AGES* females (C) or *AGES* males (D) exposed to 10 mM auxin compared to 0 mM auxin controls as determined from MetaboAnalyst 5.0 analysis.

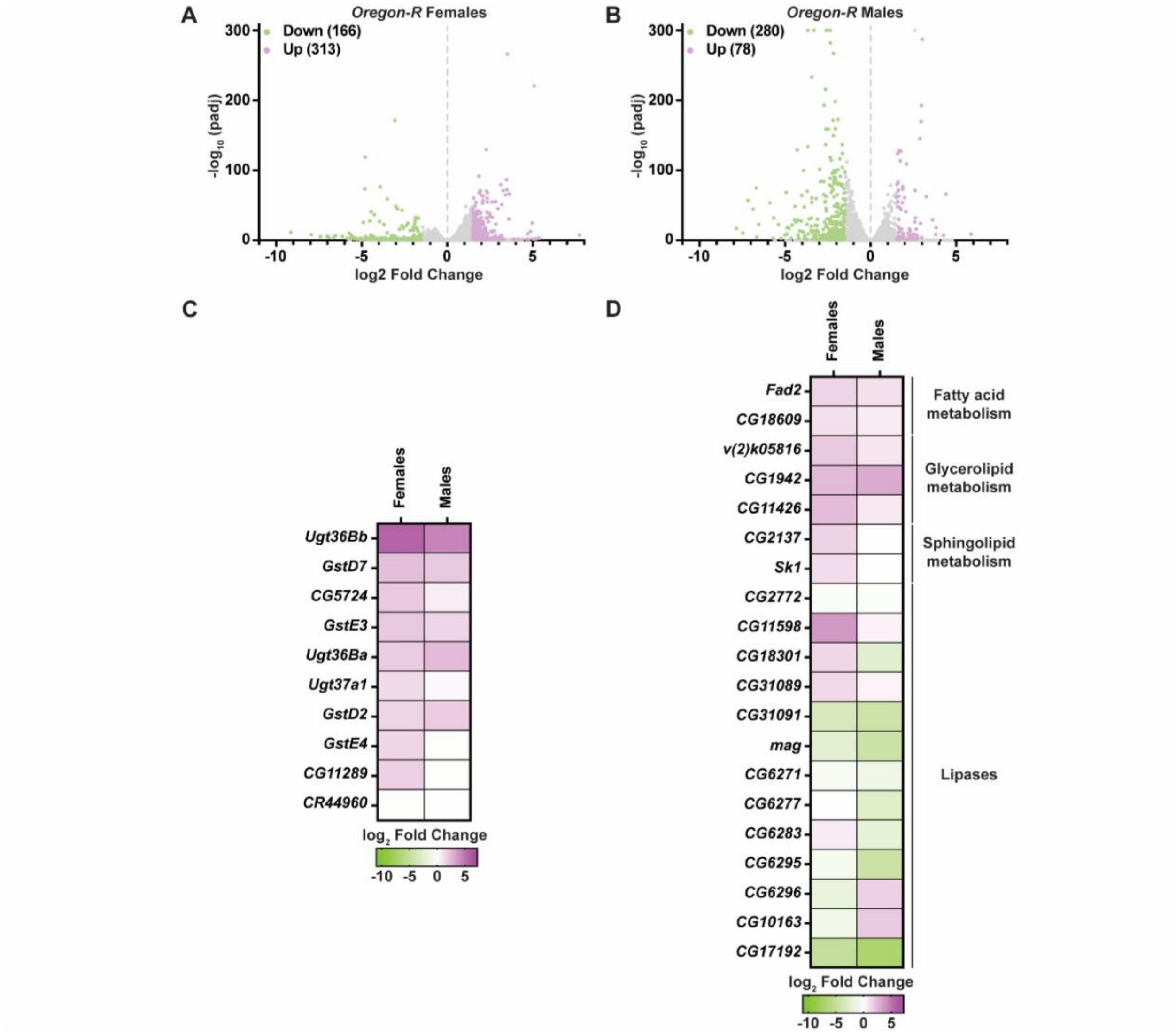


**Figure S7. Transcripts of genes that regulate fatty acid metabolism and drug detoxification are altered in *Oregon-R* males and females exposed to auxin.** **(A,B)** Volcano plots of differentially expressed genes graphing the statistical significance [-$\log_{10}$(padj)] against the magnitude of differential expression ($\log_2$ Fold Change) in *Oregon-R* females (A) or *Oregon-R* males (B) exposed to 10 mM auxin and compared to 0 mM control. **(C)** Significantly up-regulated genes involved in “drug metabolism” in adult *Oregon-R* females or males exposed to 10 mM auxin and compared to the 0 mM control. **(D)** Significantly up- (purple) and down-regulated (green)

genes involved in fatty acid metabolism in adult *Oregon-R* males or females exposed to 10 mM auxin and compared to the 0 mM control.

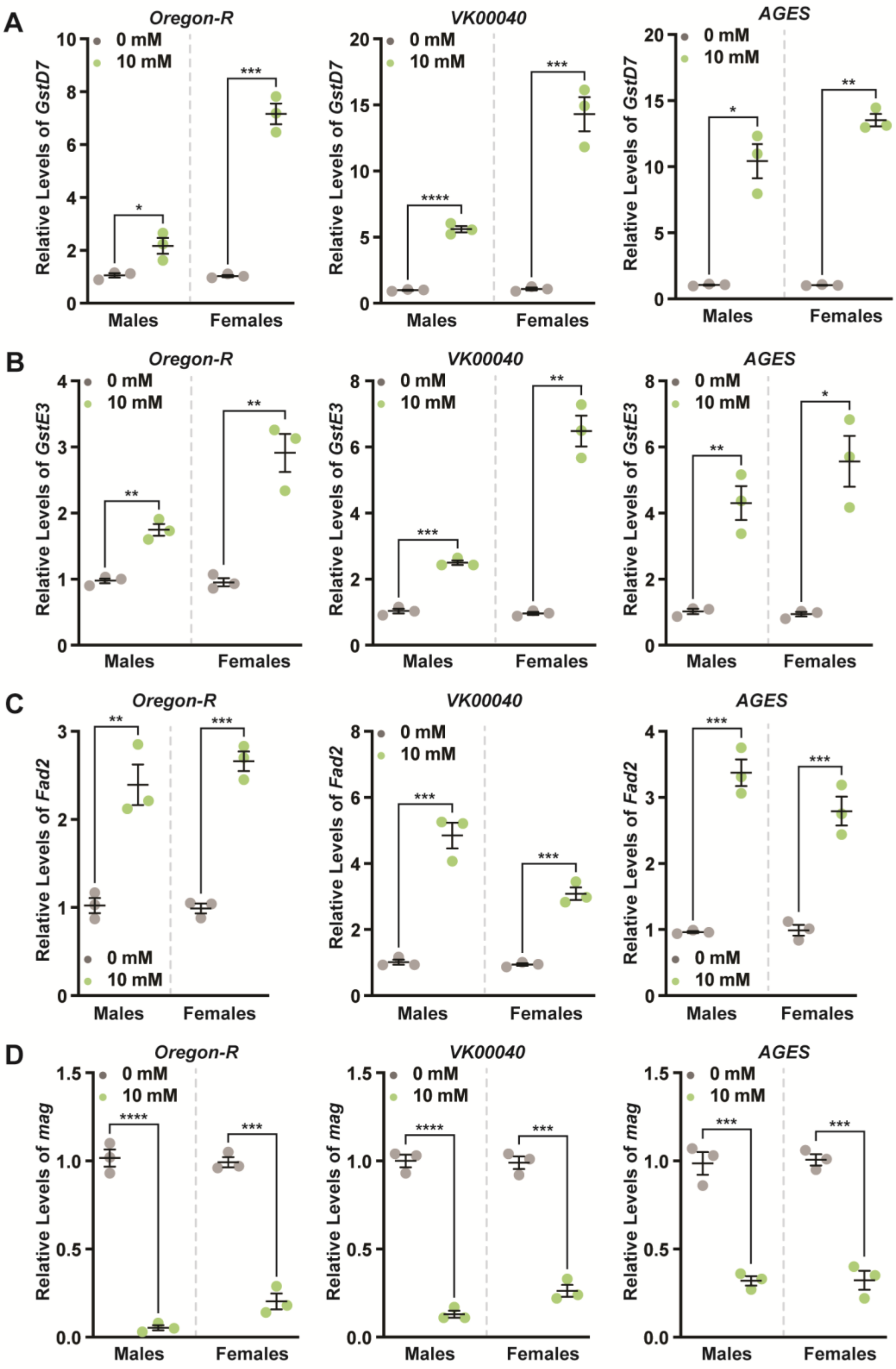
A
Oregon-R
VK00040
AGES
0 mM
10 mM
Relative Levels of GstD7
Males
Females
B
Oregon-R
VK00040
AGES
Relative Levels of GstE3
C
Oregon-R
VK00040
AGES
Relative Levels of Fad2
D
Oregon-R
VK00040
AGES
Relative Levels of mag

**Figure S8. Auxin exposure differentially regulates detoxification and fatty acid metabolism genes.** qRT-PCR analysis of *GstD7* (A), *GstE3* (B), *Fad2* (C), and *mag* (D) transcripts from whole-body males and females fed 10 mM auxin for two days compared to 0 mM control in *Oregon-R*, *VK00040*, and *AGES* backgrounds. Data shown as mean ± SEM. *$P < 0.05$, **$P < 0.01$, ***$P < 0.001$, ****$P < 0.0001$, Student's *t*-test.

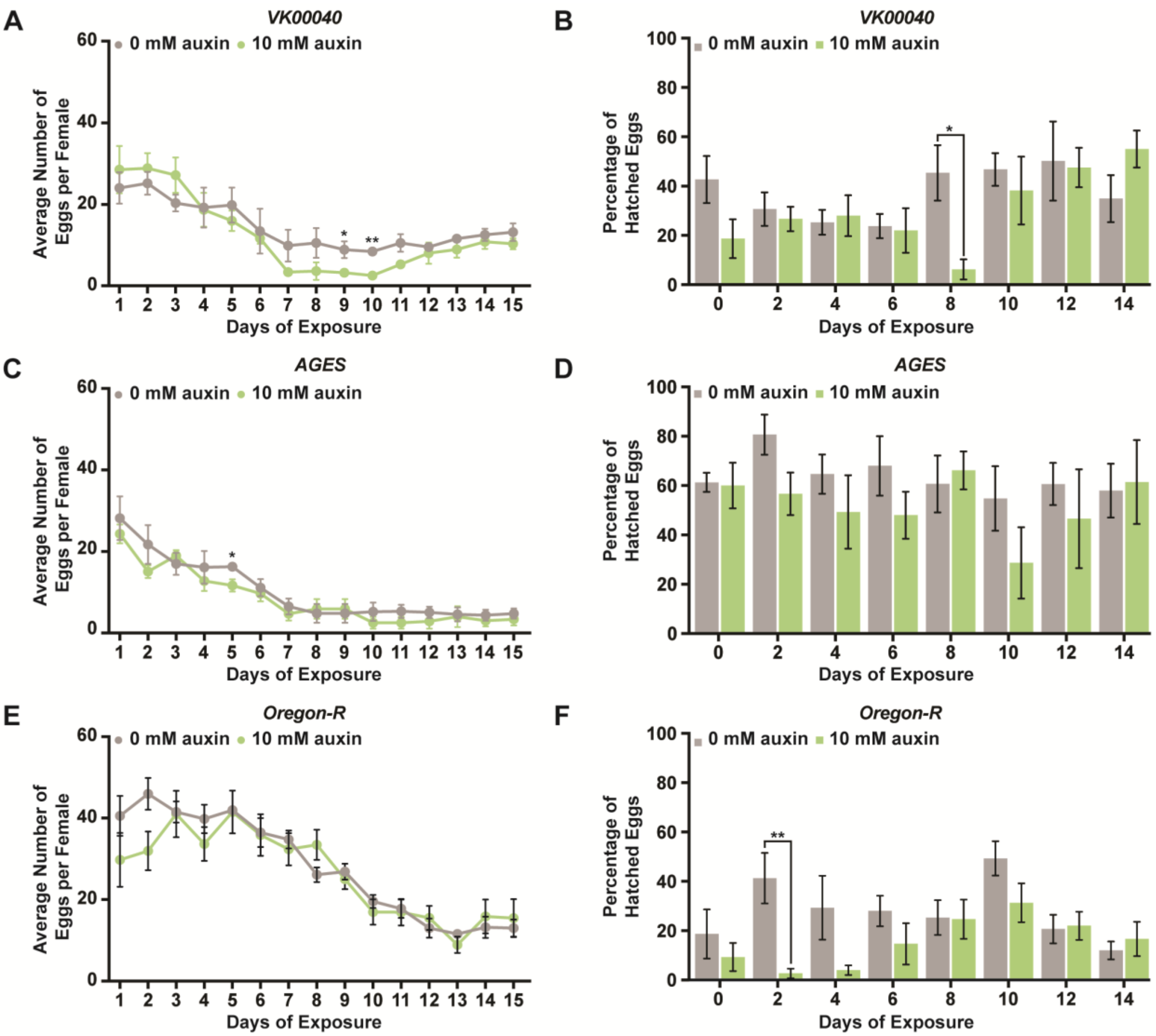


**Figure S9. Auxin exposure does not significantly alter egg laying or progeny survival.**

**(A)** Females of the *VK00040* line were incubated at $25^{o}$C and fed 0 mM or 10 mM auxin. The average number of eggs laid per female under each condition. **(B)** The average number of eggs that hatched from adult females exposed to either 0 mM or 10 mM auxin of the *VK00040* line. **(C)** Females of the *AGES* line were incubated at $25^{o}$C and fed 0 mM or 10 mM auxin. The average number of eggs laid per female under each condition. **(D)** The average number of eggs that hatched from adult females exposed to either 0 mM or 10 mM auxin of the *AGES* line. **(E)** Females of the *Oregon-R* line were incubated at 25$^{o}$C and fed 0 mM or 10 mM auxin. The average number of eggs laid per female under each condition. **(F)** The average number of eggs that hatched from adult females exposed to either 0 mM or 10 mM auxin of the *Oregon-R* line. Data is plotted as the mean ± SEM. *$P < 0.05$, **$P < 0.01$, Student's *t*-test.

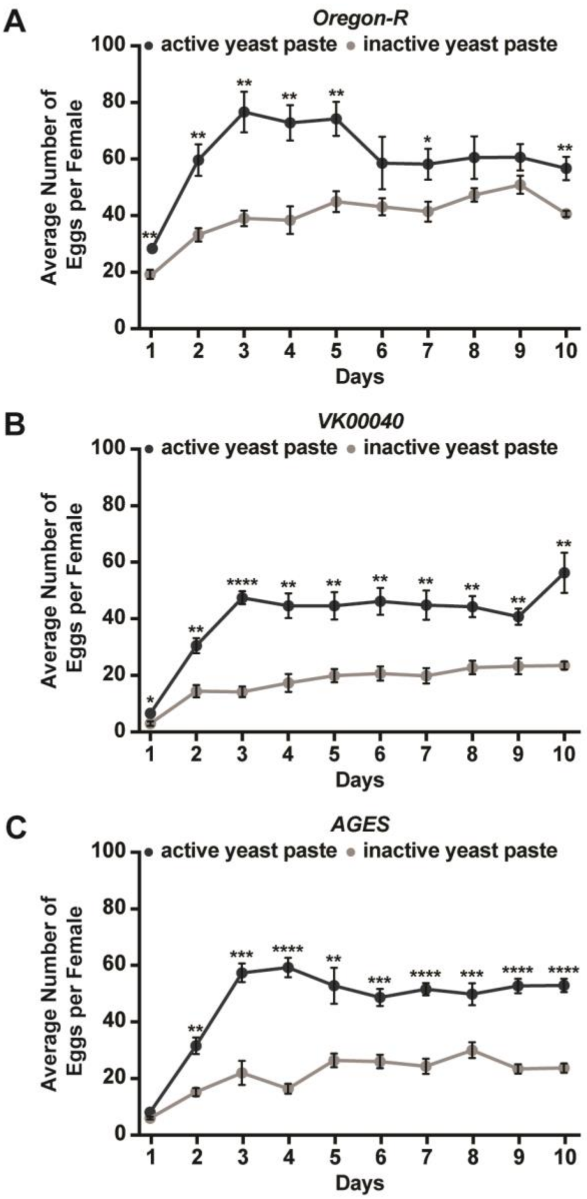


**Figure S10. Inactive yeast paste decreases egg laying in adult females.**

**(A-C)** The average number of eggs laid per female per day exposed to either active or inactive yeast paste in adult females from *Oregon-R* (A), *VK00040* (B), or *AGES* (C) backgrounds. Data shown as the mean ± SEM. $^{*}P < 0.05$, $^{**}P < 0.01$, $^{***}P < 0.001$, $^{****}P < 0.0001$, Student's *t*-test.

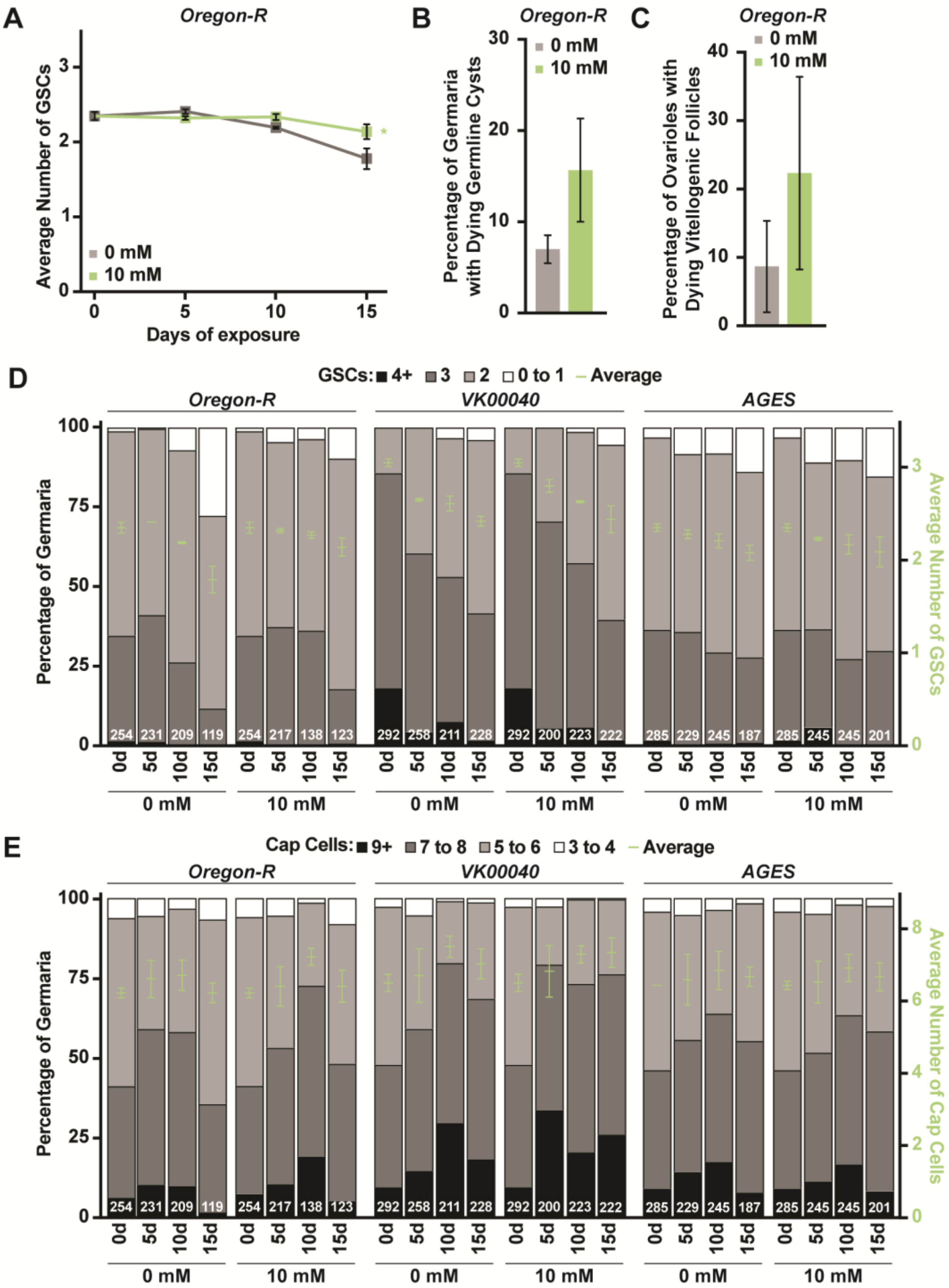


**Figure S11. Processes of oogenesis are not significantly influenced by exposure to auxin.**

**(A)** Average number of GSCs per germarium over time in *Oregon-R* females exposed to 0 mM or 10 mM auxin. Data shown as mean ± SEM. *$P < 0.05$, two-way ANOVA with interaction. **(B)** Average percentage of germaria containing ApopTag-positive germline cysts in adult *Oregon-R* females exposed to 0 mM or 10 mM auxin. Data shown as mean ± SEM; Student's $t$-test. 100 germaria were analyzed for each condition. **(C)** Average percentages of ovarioles containing dying vitellogenic egg chambers in *Oregon-R* females exposed to 0 mM or 10 mM auxin. Data shown as mean ± SEM, Student's $t$-test. 100 ovarioles were analyzed for each condition. **(D)** Bars represent the percentage of germaria containing zero, one, two, three, or four-or-more GSCs at different days of females exposed to 0 mM or 10 mM auxin in *Oregon-R*, *VK00040*, or *AGES* backgrounds. GSC number averages shown as mean ± SEM (right $y$-axis) are also plotted in Fig. 5B,C. No statistically significant differences, two-way ANOVA with interaction. Numbers of germaria analyzed are shown inside bars. **(E)** Bars represent the percentage of germaria containing three-to-four, five-to-six, seven-to-eight, ore greater or equal to nine cap cells at different times of 0 mM or 10 mM auxin exposure in *Oregon-R*, *VK00040*, or *AGES* backgrounds (left $y$-axis). Average number of cap cells per germarium shown as mean ± SEM (right $y$-axis) are plotted. No statistically significant differences, two-way ANOVA with interaction. Numbers of germaria analyzed are shown inside bars.
